# KLF4 protein stability regulated by interaction with pluripotency transcription factors overrides transcriptional control

**DOI:** 10.1101/522680

**Authors:** Navroop K Dhaliwal, Luis E Abatti, Jennifer A Mitchell

## Abstract

Embryonic stem (ES) cells are regulated by a network of transcription factors which maintain the pluripotent state. Differentiation relies on downregulation of pluripotency transcription factors disrupting this network. While investigating transcriptional regulation of the pluripotency transcription factor *Klf4*, we observed homozygous deletion of distal enhancers caused 17 fold decrease in *Klf4* transcript but surprisingly decreased protein levels by less than 2 fold indicating post-transcriptional control of KLF4 protein overrides transcriptional control. The lack of sensitivity of KLF4 to transcription is due to high protein stability (half-life >24hr). This stability is context dependent and disrupted during differentiation, evidenced by a shift to a half-life of <2hr. KLF4 protein stability is maintained through interaction with other pluripotency transcription factors (NANOG, SOX2 and STAT3) that together facilitate association of KLF4 with RNA polymerase II. In addition, the KLF4 DNA binding and transactivation domains are required for optimal KLF4 protein stability. Post-translational modification of KLF4 destabilizes the protein as cells exit the pluripotent state and mutations that prevent this destabilization also prevent differentiation. These data indicate the core pluripotency transcription factors are integrated by post-translational mechanisms to maintain the pluripotent state, and identify mutations that increase KLF4 protein stability while maintaining transcription factor function.

## Introduction

Differentiation of embryonic stem (ES) cells requires disruption to the regulatory network that maintains pluripotency gene expression. Kruppel-like factor 4 (KLF4), a member of the Kruppel-like factor family of conserved zinc finger transcription factors, is known to interact with the core network of pluripotency transcription factors (NANOG, SOX2 and OCT4) in order to regulate genes required for maintenance of pluripotency and reprogramming (Dhaliwal et al., 2018; Wei et al., 2013; Wei et al., 2009; Xie et al., 2017; Zhang et al., 2010). Sustained expression of constitutively nuclear KLF4 prevents differentiation of ES cells indicating that a loss of KLF4 protein function is required for differentiation (Dhaliwal et al., 2018). It is generally accepted that differentiation occurs due to a loss of pluripotency transcription factor activity at enhancers which disrupts pluripotency gene expression and allows for the expression of differentiation associated genes; however, the molecular mechanisms through which this disruption occurs are not well defined.

Genome-wide binding of transcription factors and coactivators can identify enhancers required for gene transcription in a particular cellular context (Chen et al., 2012; Chen et al., 2008; Moorthy et al., 2017; Visel et al., 2009). These enhancers are often located multi kb distances from the genes they regulate and form physical loops with their target gene promoters (Carter et al., 2002; Schoenfelder et al., 2015; Tolhuis et al., 2002). In pluripotent ES cells, specific enhancers have been identified that regulate *Sox2, Klf4, Nanog* and *Oct4* (*Pou5f1*) at the transcriptional level (Blinka et al., 2016; Li et al., 2014; Xie et al., 2017; Yeom et al., 1996; Zhou et al., 2014). In addition to these intrinsic mechanisms that regulate the expression of pluripotency transcription factors, the balance between maintaining the pluripotent state and inducing differentiation is also modulated by cell extrinsic factors and cell signaling cascades. Leukemia inhibitory factor (LIF) maintains the pluripotent state by activating JAK-STAT signaling causing phosphorylation and activation of STAT3 (Matsuda et al., 1999; Niwa et al., 1998). Activated STAT3 induces transcription of *Klf4* through binding to the enhancers downstream of *Klf4* (Hall et al., 2009; Xie et al., 2017; Zhang et al., 2010). In addition, dual inhibition (2i, GSK3 and MEK inhibition) maintains ES cells in a naïve state closest to that of the precursor cells from the pluripotent epiblast of pre-implantation embryos (Nichols and Smith, 2009; Tosolini and Jouneau, 2016; Wray et al., 2010).

As pluripotency master regulators are transcription factors, reduced transcription of specific genes is generally considered the mechanism through which differentiation of ES cells occurs, however, changes in gene transcription do not always correlate with changes in protein levels. At a genome scale, evaluation of the correlation between mRNA abundance and protein abundance estimates that, for cells in a steady state, 50-80% of the variability in protein levels can be explained by the levels of mRNA present (reviewed in (Liu et al., 2016). For cells undergoing dynamic transitions, for example during monocyte to macrophage differentiation, protein and mRNA levels become decoupled during the early differentiation phase, mainly due to a delay in translation compared to transcription (Kristensen et al., 2013). In both cases exceptions exist where mRNA and protein levels do not correlate even when delays in translation are taken into account, however, the mechanism through which this occurs is not well understond. Transcription factors generally display low protein stability which allows rapid cell state transitions (Hochstrasser and Varshavsky, 1990; Jovanovic et al., 2015; Zhou et al., 2004). In this study, however, we show that KLF4 protein levels are highly decoupled from the RNA levels due to exceptional stability of the KLF4 protein in naïve ES cells maintained in LIF/2i. Homozygous deletion of downstream *Klf4* enhancer regions caused a 17 fold reduction in *Klf4* transcript levels whereas KLF4 protein levels were reduced by <2 fold. Surprisingly, we observed a greater reduction of KLF4 protein levels (>3 fold) in ES cells with compromised SOX2 expression, despite the observation that *Klf4* transcript levels are unchanged in these cells. We found that these discrepancies in KLF4 protein and transcript levels are due to modulation of KLF4 protein stability by SOX2, NANOG and activated STAT3 as well as domains within the KLF4 protein that anchor KLF4 in the nucleus. During pluripotency exit KLF4 protein becomes destabilized. Preventing this destabilization through mutation of KLF4 destabilizing motifs blocks pluripotency exit. The core pluripotency maintenance transcription factors are known to function in a highly integrated way to maintain transcriptional control of the pluripotent state. Here we show a new way in which these factors regulate each other that bypasses transcriptional control but maintains post-translational control of KLF4 function.

## Results

### *Klf4* transcript and protein levels are uncoupled in embryonic stem cells maintained in LIF/2i

For ES cells maintained in LIF/serum, *Klf4* has been shown to be regulated by three enhancers 54-68kb downstream of the gene; deletion of this region was found to reduce *Klf4* transcription by 90%, greatly affecting KLF4 protein levels (Xie et al., 2017). For ES cells maintained in the more naïve state by LIF/2i, we determined that although the enhancers remain important for maintaining transcript levels, functional KLF4 protein is maintained in the absence of the enhancers. We used F1 (*Mus musculus*^129^ x *Mus castaneus*) ES cells, allowing allele-specific deletion screening and gene expression analysis (Moorthy and Mitchell, 2016; Zhou et al., 2014). Upon deletion of two (Δ1, 8 fold reduction in RNA), or all three enhancers (Δ2, 17 fold reduction in RNA) we observed a dramatic reduction in *Klf4* transcript levels but a much more subtle change in KLF4 protein levels (Figure 1 and S1). KLF4 protein levels are significantly reduced only in cells with the Δ2 homozygous deletion (Δ2^129/Cast^) and in these cells, which displayed a 17 fold reduction in mRNA, protein was reduced by <2 fold. To confirm this was not an effect of recent enhancer deletion, we investigated *Klf4* transcript and protein levels in cells maintained to later passages (P9), but identified no significant differences between early and late passages. Transcript and protein levels of other pluripotency transcription factors, *Oct4, Sox2, Nanog, Klf2*, and *Klf5* remained unchanged in *Klf4* enhancer deleted clones (Figure 1 and S1).

**Figure 1:**
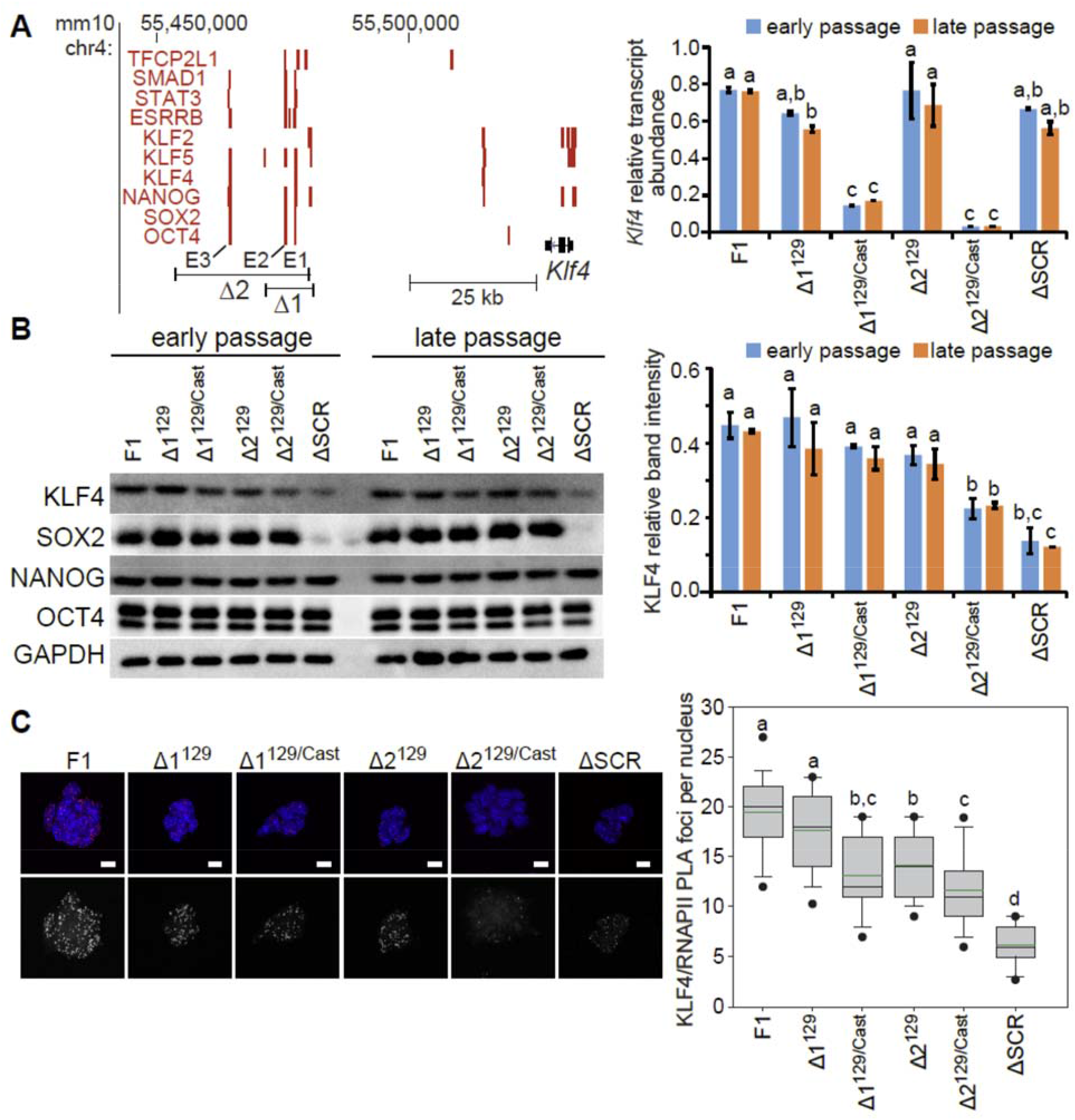
*Klf4* transcript and protein levels are uncoupled in embryonic stem cells maintained in LIF/2i. A) Genome browser view (left) of enhancer regions (E1, E2, E3) downstream of *Klf4*, indicating Δ1 and Δ2 deletions. Red bars show binding of transcription factors involved in pluripotency. Data are displayed on the mm10 assembly of the University of California at Santa Cruz (UCSC) Genome Browser. Total *Klf4* transcript levels (right) quantified relative to *Gapdh* from three biological replicates of both early and late passages of *Klf4* enhancer deleted clones and the ΔSCR^129/Cast^ clone (ΔSCR). Error bars represent standard deviation, statistical differences determined by two way ANOVA (P < 0.05) are indicated by different letters. B) Immunoblots (left) showing the levels of pluripotency transcription factors in both early and late passages for *Klf4* enhancer deleted clones. GAPDH levels indicate sample loading. KLF4 quantified relative to GAPDH (right) from three biological replicates of early and late passages for *Klf4* enhancer deleted clones and the ΔSCR^129/Cast^ clone (ΔSCR). Error bars represent standard deviation, statistical differences determined by two way ANOVA (P < 0.05) are indicated by different letters. C) Proximity ligation amplification (PLA) displays the interaction between KLF4 and RNAPII in *Klf4* enhancer deleted and ΔSCR^129/Cast^ clones (ΔSCR). Images shown are maximum-intensity projections. Merged images at the top display DAPI in blue and PLA in red; PLA signal shown in grey scale at the bottom. Scale bar = 10 μm. On the right box-and-whisker plots indicate the number of PLA foci per nucleus. Boxes indicate interquartile range of intensity values and whiskers indicate the 10th and 90th percentiles; outliers are shown as black dots. Images were collected from at least three biological replicates and ≥100 nuclei were quantified for each clone. Statistical differences determined by one-way ANOVA (P < 0.05) are indicated by different letters.

Surprisingly, we observed that KLF4 protein levels were significantly reduced by >3 fold in clones with reduced SOX2 protein levels due to a homozygous deletion of the enhancer that regulates *Sox2* transcription in ES cells (ΔSCR^129/Cast^)(Li et al., 2014; Zhou et al., 2014), despite the observation that *Klf4* transcript levels are unaffected by SCR deletion (Figure 1A). To examine the levels of functional nuclear KLF4 protein, we used proximity ligation amplification (PLA) to investigate the interaction between KLF4 and active serine 5 phosphorylated RNAPII (RNAPII-S5P) in individual nuclei. Similar to the results for total KLF4 protein levels, we found only a subtle reduction in KLF4/RNAPII interaction in cells with the KLF4 enhancer deletions; however, we observed that the greatest reduction in KLF4 association with RNAPII occurred in cells with reduced SOX2 protein levels (ΔSCR^129/Cast^, Figure 1C). Together these data reveal a disconnect between the levels of *Klf4* RNA and protein in naïve ES cells and indicate the KLF4 protein may be highly stable in the naïve pluripotent state.

### KLF4 protein stability is regulated by LIF and MAPK signaling pathways

Investigation of KLF4 protein stability in naïve ES cells maintained in LIF/2i and cells differentiated for 24hr revealed that KLF4 protein is more stable in undifferentiated cells with a t_½_ >24hr. After removal of LIF/2i for 24hr, KLF4 becomes unstable with a t_½_ <2 hr (Figure 2A). By contrast, the other pluripotency transcription factors, OCT4, SOX2 and NANOG, are unstable (t_½_ 2-4hr) in undifferentiated cells and their stability was not affected by differentiation (Figure S2). Similarily the other ES cell-expressed *Klf* family proteins, KLF2 and KLF5, are not as stable as KLF4 with t_½_ ~3hr and their stability is not affected by differentiation (Figure S2). KLF4 protein stability is not affected by deletion of its downstream enhancers, explaining the modest reduction in KLF4 protein in these clones (Figure S2).

**Figure 2:**
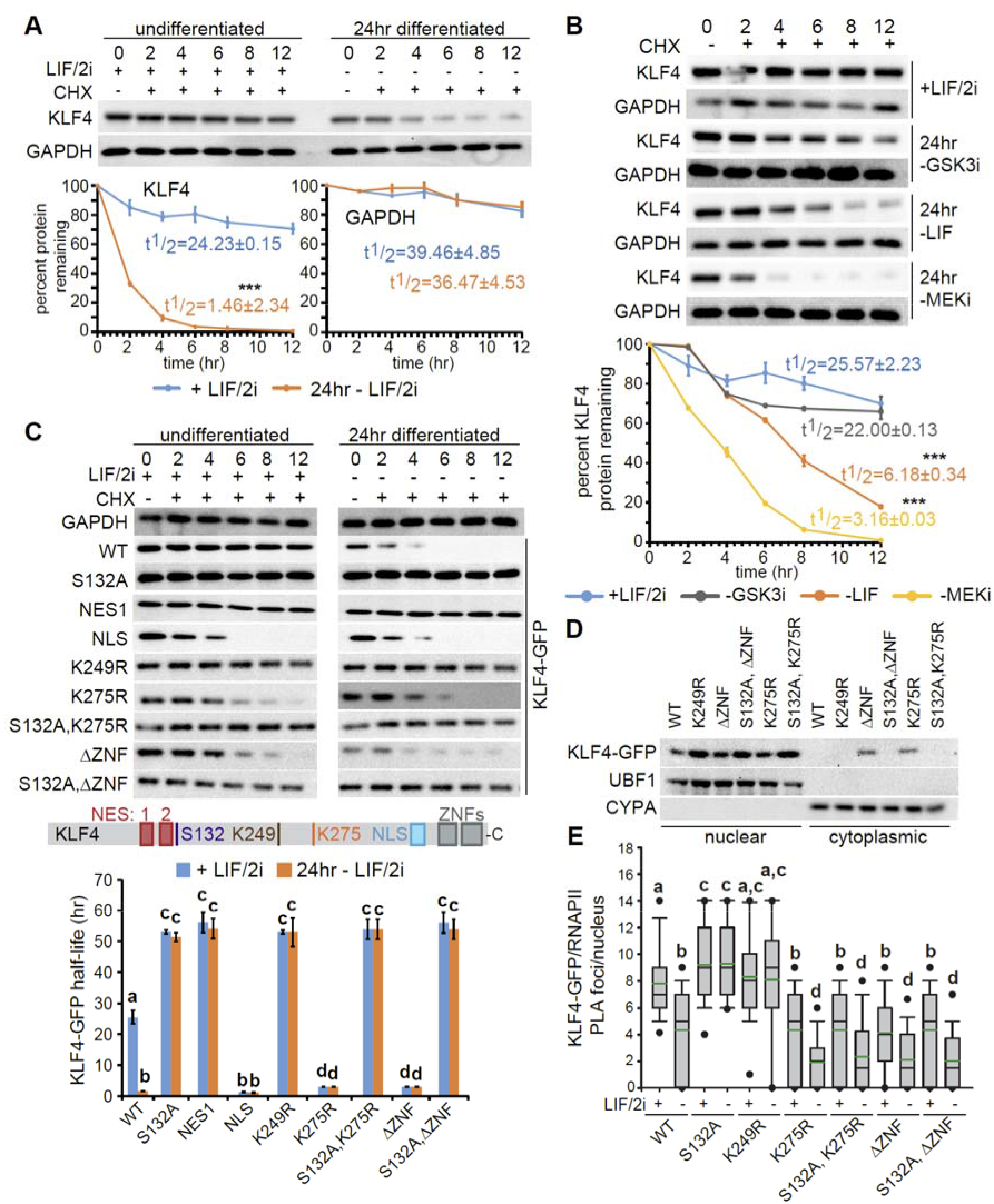
KLF4 protein stability is regulated by the LIF and MAPK signaling pathways, nuclear anchoring and post-translational modifications to KLF4. A) Immunoblots for KLF4 and GAPDH in ES cells cultured with LIF/2i (undifferentiated) and 24hr after removal of LIF/2i, sampled at 0, 2, 4, 6, 8 and 12hr after CHX treatment. GAPDH levels were used as a control and displayed the expected half-life (t_½_ >30hr). On the bottom percent remaining KLF4 or GAPDH protein was calculated from the intensity of CHX treatment immunoblots, measured in three biological replicates. KLF4 or GAPDH protein half-life in the indicated clones maintained in the presence or absence of LIF/2i was calculated for each time series replicate by best fit to exponential decay. Error bars represent standard deviation of three biological replicates. Statistical differences determined by two-tailed t-test are indicated by *** (P < 0.001). B) Immunoblots for KLF4 and GAPDH in ES cells cultured with LIF/2i and 24hr after removal of individual media components (GSK3i, LIF, or MEKi), sampled at 0, 2, 4, 6, 8 and 12hr after CHX treatment. On the bottom percent remaining KLF4 protein was calculated from the intensity of CHX treatment immunoblots, measured in three biological replicates. Half-life was calculated for each time series replicate by best fit to exponential decay. The error bars represent standard deviation. Statistical differences between protein half-life in different culture conditions compared to LIF/2i were determined by two-tailed t-test (P < 0.001) and are indicated by ***. C) Immunoblots for WT KLF4-GFP, the indicated mutants, and GAPDH from ES cells cultured with LIF/2i and 24hr after LIF/2i removal, sampled at 0, 2, 4, 6, 8 and 12hr after CHX treatment. In the middle a schematic of mouse KLF4 is shown depicting nuclear export signals (NES), ERK phosphorylation site S132, ubiquitination site K249, sumoylation site K275, nuclear localization signal (NLS) and zinc fingers (ZNFs). At the bottom the calculated KLF4 protein half-life is shown in the indicated mutants cultured with LIF/2i and 24hr after LIF/2i removal. Half-life was calculated for each time series replicate by best fit to exponential decay. Error bars represent standard deviation of three biological replicates. Statistical differences determined by two-way ANOVA (P < 0.05) are indicated by different letters. D) Nuclear and cytoplasmic fractions prepared from WT KLF4-GFP and the indicated mutants, immunoblots probed with anti-GFP, indicated the expression and localization of KLF4-GFP. UBF1 and CYPA were used to analyze the purity of nuclear and cytoplasmic fractions respectively. E) Box-and-whisker plots display the number of KLF4-GFP/RNAPII proximity ligation amplification (PLA) foci per nucleus for WT and the indicated mutants. Boxes indicate interquartile range of intensity values and whiskers indicate the 10th and 90th percentiles; outliers are shown as black dots. Images were collected from at least three biological replicates and ≥100 nuclei were quantified for each sample. Statistical differences determined by two way ANOVA (P < 0.05) are indicated by different letters.

Reduced KLF4 protein stability upon differentiation by removal of LIF/2i suggested extrinsic factors that activate signaling cascades could be involved in regulating KLF4 protein stability. Upon investigating the effect of individual signaling pathways on KLF4 stability, we observed that activation of the MAPK pathway, which occurs after MEKi removal, has the most significant effect on KLF4 stability, followed by inhibition of the JAK-STAT pathway by LIF removal (Figure 2B). Removing GSK3i, allowing activation of Wnt signaling for 24hr, did not have a significant effect on KLF4 stability. For ES cells maintained in LIF/serum, KLF4 displayed low protein stability similar to the observed stability after removal of MEKi (Figure S2). Previous studies demonstrated that these signaling pathways regulate gene transcription (Dhaliwal et al., 2018; Nichols and Smith, 2009; Theunissen et al., 2011; Zhang et al., 2010); therefore, we investigated the effect of individual signaling pathways on *Klf4, Nanog, Oct4, Sox2, Klf2* and *Klf5* transcription. We observed that the removal of MEKi alone or in combination with LIF/GSK3i for 12hr significantly reduces *Klf4* and *Nanog* transcription, but *Oct4, Sox2, Klf2* and *Klf5* transcription remains unaffected (Figure S3). As changes to *Klf4* transcription could confound the analysis of KLF4 stability during differentiation, we investigated the stability of KLF4-GFP controlled by a CMV promoter upon removal of MEKi, LIF or GSK3i (Figure S4). The stability of KLF-GFP in these conditions was similar to the stability of the endogenous protein, with removal of MEKi or LIF reducing KLF4 stability more dramatically than removal of GSK3i.

The MAPK pathway has been shown to have a role in KLF4 nuclear localization, ubiquitination and degradation (Dhaliwal et al., 2018; Kim et al., 2012; Kim et al., 2014). As early as 6hr after MEKi removal, KLF4 was observed to exit the nucleus, and cytoplasmic KLF4 undergoes proteosomal degradation (Dhaliwal et al., 2018). In addition, nuclear export was found to depend on the presence of both a KLF4 nuclear export signal (NES1 at 97-107) and the ERK phosphorylation site in KLF4 at S132 (Dhaliwal et al., 2018). KLF4 nuclear export after MEKi removal, allowing ERK activation, could explain the observed reduction in KLF4 protein stability; to investigate this further we used ES cells with stable integration of wild-type (WT) KLF4-GFP, the NES1 mutant (KLF4(NES1)-GFP), KLF4(S132A)-GFP and a nuclear localization sequence (NLS) mutant, KLF4(NLS)-GFP. WT KLF4-GFP displayed stability similar to endogenous KLF4 with a t_½_ >24hr in undifferentiated cells which was reduced to <2hr after 24hr differentiation (Figure 2C). Both of the constitutively nuclear mutants, KLF4(NES1)-GFP and KLF4(S132A)-GFP, were highly stable proteins (t_½_ >51hr) and this stability was not affected by differentiation. By contrast, the constitutively cytoplasmic KLF4(NLS)-GFP was unstable, with a t_½_ <2hr in both undifferentiated and differentiated cells. Together these data indicate nuclear localization is critical for KLF4 protein stability.

### KLF4 association with DNA and RNAPII is required to maintain protein stability through nuclear anchoring

As localization to the nucleus increased the stability of KLF4, we next investigated protein stability of mutants with disrupted KLF4 transcription factor function. KLF4 contains two c-terminal zinc finger (ZNF) domains required for DNA binding (Schuetz et al., 2011; Wei et al., 2009) and a sumoylation site at K275 shown to be important for transactivation of target promoters in reporter assays (Du et al., 2010). The amino acids surrounding K275 are highly conserved highlighting conservation of the KLF4 sumoylation site (Figure S5). Stable expression of KLF4 loss of function mutants, with a mutated sumoylation site (KLF4(K275R)-GFP) or deleted zinc fingers (KLF4ΔZNF-GFP), were disrupted in their nuclear localization (Figure 2D) and were more unstable compared to WT KLF4-GFP protein with a t_½_ of ~3hr (Figure 2C). Mutation of S132, together with K275 or zinc finger deletion, restored nuclear anchoring and increased the stability of KLF4 (Figure 2C/D). Interestingly, disruption of either KLF4 sumoylation (K275R) or DNA binding (ΔZNF) interfered with recruitment to RNAPII-S5P nuclear compartments, as shown by the decrease in the number of KLF4/RNAPII PLA foci per nucleus compared to WT KLF4-GFP (Figure 2E). Introducing the S132 mutation into the KLF4 loss of function mutants restored nuclear anchoring but did not restore RNAPII association, indicating that sumoylation and the DNA binding domains are required for KLF4 association with RNAPII.

### Ubiquitination of KLF4 is required for nuclear export and degradation during the course of differentiation

Upon phosphorylation by ERK and subsequent nuclear export, KLF4 has been shown to be degraded causing ES cell differentiation (Dhaliwal et al., 2018; Kim et al., 2012). As expected, treatment with the proteasome inhibitor (MG132) prevented the decrease in the levels of KLF4 protein normally observed in cells differentiated for 24hr (Figure 3A). A previous study showed that K232 was the most critical residue in human KLF4 for ubiquitination and degradation (Lim et al., 2014). KLF4 is highly conserved in this region with K249 in the mouse sequence predicted as a ubiquitination site by UbPred and NetChop (Figure S5)(Kesmir et al., 2002; Radivojac et al., 2010). In order to further investigate the role of K249 in KLF4 function, we generated stable ES lines expressing a KLF4 ubiquitination site mutant, KLF4(K249R)-GFP. KLF4(K249R)-GFP is nuclear in undifferentiated ES cells, similar to the WT protein (Figure 2D), but displayed an increased t_½_ of 53hr independent of culture conditions similar to the S132 and NES1 mutants (Figure 2C), indicating that blocking KLF4 ubiquitination prevents the loss of KLF4 stability upon differentiation. Blocking KLF4 ubiquitination by mutation of K249 does not disrupt interaction with RNAPII-S5P in undifferentiated cells and prevents the loss of KLF4/RNAPII-S5P interaction associated with differentiation (Figure 2E). As this was similar to what occurs in the KLF4 S132 and NES1 mutants, which showed disrupted nuclear export and no interaction with Xportin 1 (XPO1) (Dhaliwal et al., 2018), we investigated the interaction between KLF4(K249R)-GFP and XPO1 in differentiating ES cells by PLA. Indeed mutation of K249 did disrupt the interaction between KLF4 and XPO1 normally observed in differentiating cells (Figure 3B).

**Figure 3:**
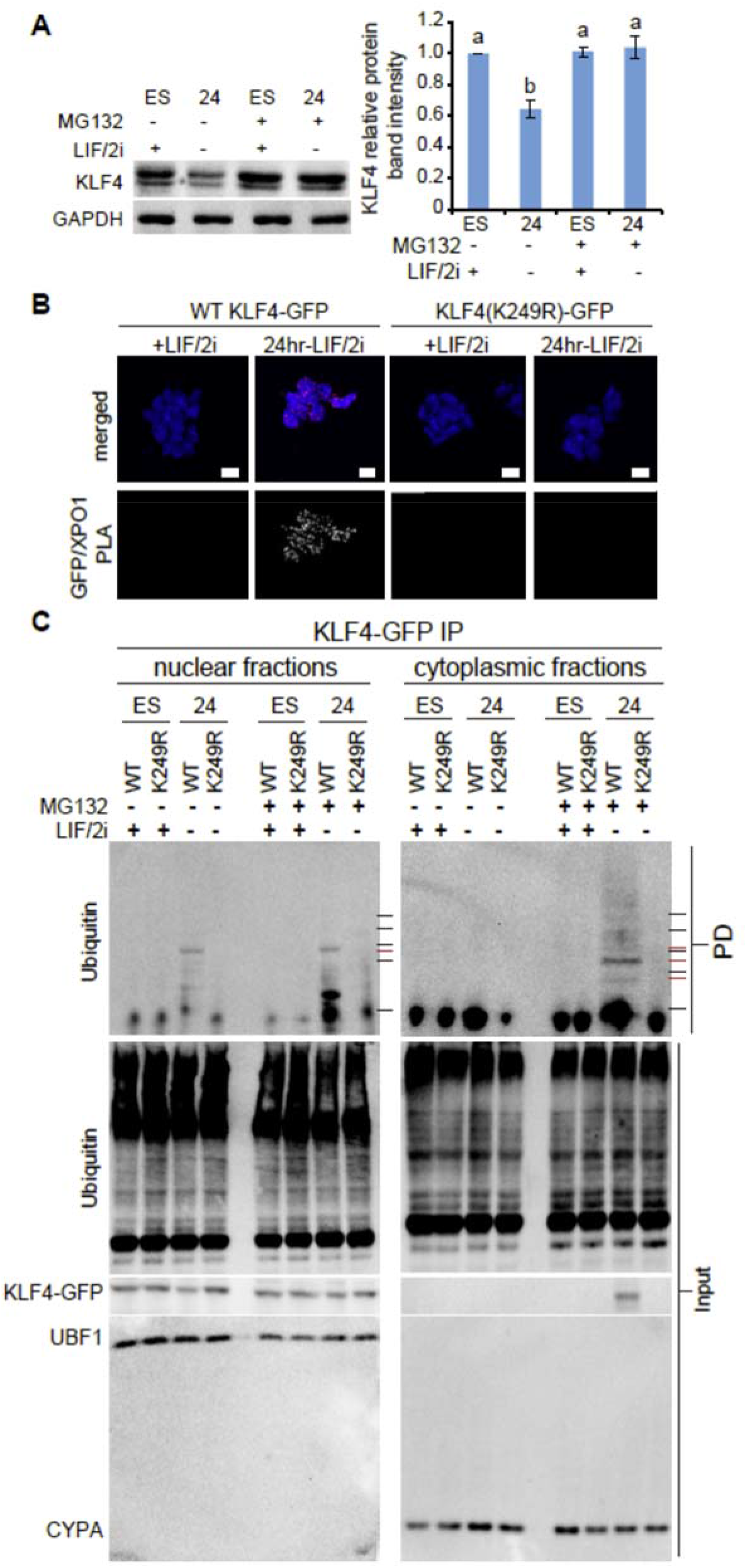
Ubiquitination of KLF4 is required for nuclear export and degradation during differentiation. A) Immunoblots for KLF4 in whole cell lysate prepared from ES cells cultured with LIF/2i, and 24hr after removal of LIF/2i, treated or not with proteasome inhibitor MG132. GAPDH levels indicate sample loading. On the right quantification of KLF4 protein intensity level relative to GAPDH and ES cell levels (+LIF/2i, - MG132), in three biological replicates. Error bars represent standard deviation. Statistical differences determined by one way ANOVA (P < 0.05) are displayed by different letters. B) Proximity ligation amplification (PLA) displays the interaction between KLF4-GFP/XPO1 in WT and K249R mutant expressing cells cultured with LIF/2i and 24hr after removal of LIF/2i. Images shown are maximum-intensity projections. Merged images display DAPI in blue and PLA in red. Scale bar =10 μm. C) KLF4-GFP immunoprecipitation (IP) using anti-GFP from nuclear and cytoplasmic fractions of WT and the K249R mutant expressing cells cultured with LIF/2i and 24hr after removal of LIF/2i, treated and untreated with proteosomal inhibitor MG132. Input and pull-down (PD) samples were probed with anti-Ubiquitin. To the right of the Ubiquitin blot molecular weight markers are indicated by black lines; from the top 250, 150, 100, 75, 50 kDa. The red lines indicate bands with calculated molecular weights of 95 kDa in the nuclear fraction and 105, 84 and 72 kDa in the cytoplasmic fraction. Cyclophilin A (CYPA) and the nucleolar protein upstream binding factor (UBF1) were detected simultaneously for all samples and reveal purity of the cytoplasmic and nuclear fractions, respectively.

We next investigated the ubiquitination status of KLF4 in nuclear and cytoplasmic fractions and the effect of K249 mutation on KLF4 ubiquitination (Figure 3C). KLF4-GFP protein has a predicted molecular weight of 81 kDa; in ES cells immunoblot for KLF4-GFP protein identifies a prominent band at 84 kDa (Figure 3C, input). WT KLF4-GFP immunoprecipitated from the nuclear fraction using an anti-GFP antibody displayed a prominent ubiquitin band at 95 kDa (Figure 3C), which could correspond to monoubiquitation of KLF4-GFP as ubiquitin has a molecular weight of 8.5 kDa. This band was only apparent after removal of LIF/2i and was absent from the KLF4(K249R)-GFP mutant, revealing that ubiquitination depends on the presence of K249. WT KLF4-GFP immunoprecipitated from cytoplasmic fractions using an anti-GFP antibody revealed more of a high molecular weight laddering pattern above 105 kDa which could correspond to polyubiquitination of cytoplasmic KLF4-GFP in MG132-treated, differentiated (-LIF/2i) cells. The presence of bands below 95 kDa could represent partial KLF4-GFP degradation (Figure 3C). These ubiquitin immunoreactive bands were not detected in the absence of MG132 suggesting cytoplasmic polyubiquitinated KLF4 is rapidly degraded. KLF4(K249R)-GFP was not detected in the cytoplasmic fraction by either ubiquitin or GFP antibodies, indicating that mutation of the ubiquitination site blocked nuclear export of KLF4-GFP.

### The loss of KLF4 protein stability is required for ES cell differentiation

To evaluate the role of KLF4 stability in pluripotency maintenance and exit from the pluripotent state, we differentiated cells expressing WT KLF4-GFP or mutant proteins for 5 days in the absence of LIF/2i. Cells in the pluripotent state exhibit high alkaline phosphatase activity which is lost upon differentiation (Štefková et al., 2015). Cells that express KLF4 mutants with increased protein stability [KLF4(K249R)-GFP and KLF4(S132A)-GFP] maintain alkaline phosphatase activity 5 days after removal of LIF/2i, indicating a block in differentiation (Figure 4A). In addition, expression of either KLF4(K249R)-GFP or KLF4(S132A)-GFP prevented decreased expression of endogenous pluripotency transcription factors normally downregulated during differentiation after removal of LIF/2i (Figure 4B).

**Figure 4:**
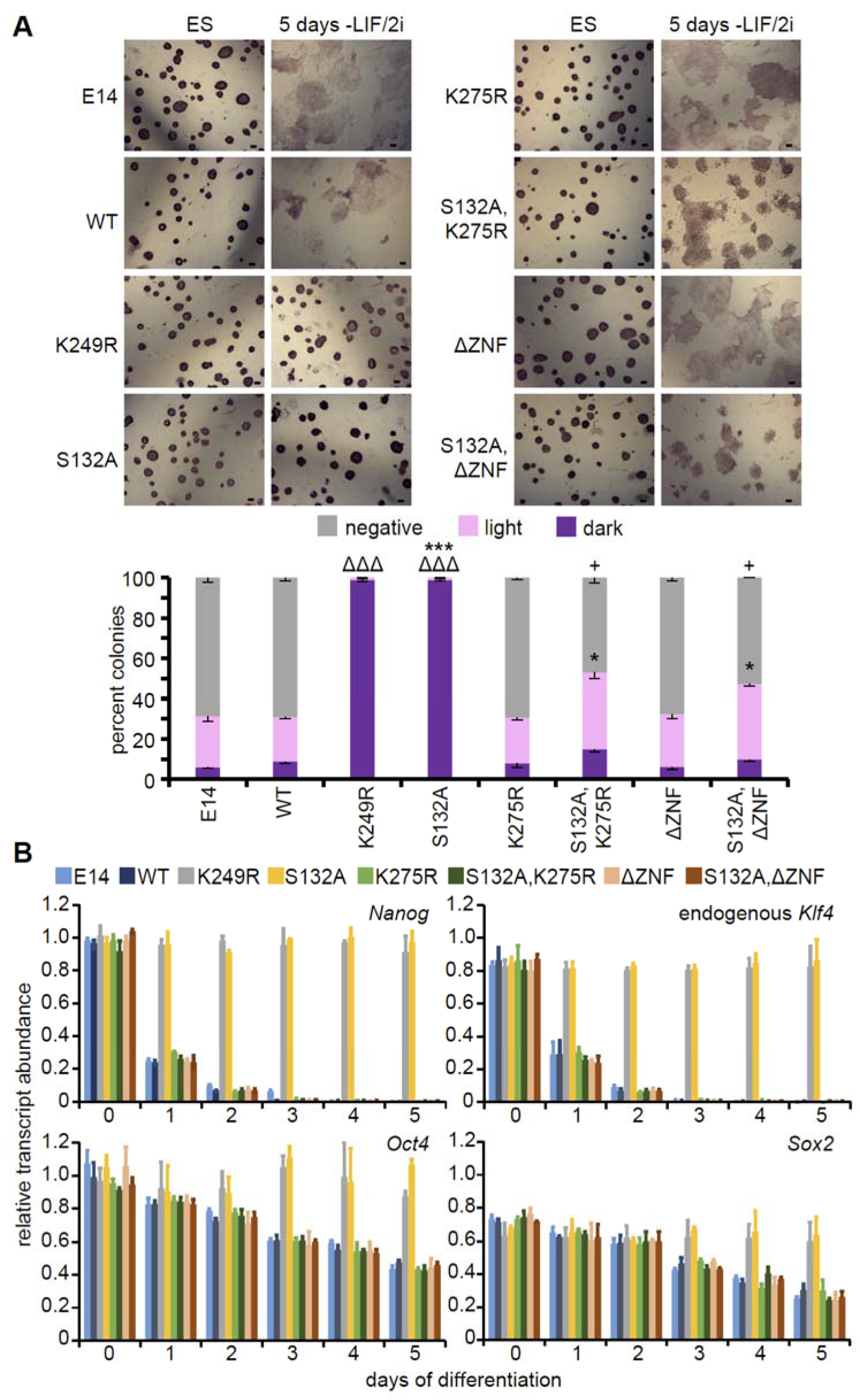
Loss of KLF4 protein stability is required for ES cell differentiation. A) Alkaline phosphatase staining of untransfected E14 ES cells, WT and the indicated KLF4-GFP mutant expressing cells in undifferentiated ES cells and 5 days after LIF/2i removal. Scale bar = 50μm. After 5 days of LIF/2i removal positive and negative colonies were counted from at least three replicates for each indicated mutant, revealing that expression of WT KLF4-GFP does not block differentiation whereas expression of the S132A or K249R mutant does. Error bars represent standard deviation. Statistical differences between each type of colony compared to WT KLF4-GFP determined by t test are indicated by ΔΔΔ P < 0.001 for dark colonies, * P < 0.5, *** P < 0.001 for light colonies and + P < 0.5 for negative colonies. B) Transcripts for endogenous *Klf4, Nanog, Oct4* and *Sox2* were quantified relative to *Gapdh* levels, in three biological replicates of untransfected E14 ES cells, WT and the indicated KLF4-GFP mutant cells revealing that only KLF4(S132A)-GFP, KLF4(K249R)-GFP blocked differentiation. Error bars represent standard deviation.

To further investigate the importance of the role of KLF4 as a transcription factor in this context, we evaluated the role of the sumoylation site that we found was important for recruitment to the RNAPII compartment and which has been shown to be involved in transactivation (Du et al., 2010). Although expression of the K275 sumoylation site mutant does not affect differentiation, mutation of this site in an S132A ERK phosphorylation site mutant background abolished the ability of the compound mutant to maintain naïve pluripotency (Figure 4). Similarly, we found that KLF4 requires the ability to bind DNA through its zinc finger domains, as differentiation occurred in cells expressing the compound mutant, KLF4(S132AΔZNF)-GFP, indicating this mutant was unable to maintain pluripotency in the absence of LIF/2i. Together these data indicate that maintaining high KLF4 protein stability blocks pluripotency exit and KLF4 requires the ability to bind DNA and associate with RNAPII to maintain the pluripotent state.

### Phosphorylated STAT3 stabilizes KLF4 protein in nuclear complexes

Removal of LIF from cells maintained in LIF/2i significantly reduced KLF4 protein stability from t_½_ = 26hr to t_½_ = 6hr after 24hr (Figure 2B). LIF signaling activates the JAK-STAT pathway in ES cells causing phosphorylation and activation of STAT3 (Hirai et al., 2011; Ohtsuka et al., 2015; Raz et al., 1999; Zhang et al., 2010). After removal of LIF/2i the levels of phosphorylated STAT3 (pSTAT3) in the nucleus decrease rapidly and are undetectable after 6hr of differentiation (Figure S6). PLA for STAT3/KLF4 during early differentiation showed that interaction between these two proteins occurs in undifferentiated cells but is lost during the first 24hr of differentiation (Figure S6). We next examined whether a short 1hr LIF induction in cells differentiated for 24hr could restore STAT3/KLF4 interaction and KLF4 protein stability. The t_½_ of KLF4 increased form <2hr to 24hr after a 1hr treatment with LIF (Figure 5A). This short LIF treatment also increased the levels of *Klf4* transcript and protein in 24hr differentiated cells (Figure 4D/E), and restored the interaction between KLF4 and STAT3 (Figure 5B-D). As LIF regulated KLF4 at both the transcriptional and post-transcriptional levels, we repeated the 1hr LIF treatment in cells treated at the same time with the protein synthesis inhibitor cycloheximide (CHX). Even in the presence of CHX, inhibiting de novo protein synthesis, an increase in KLF4 stability was observed after 1hr LIF treatment (Figure S7). Next, we investigated KLF4 association with RNAPII by PLA which revealed a significant increase in the number of interaction foci per nucleus after the 1hr LIF induction (Figure 5E). These data indicate that pSTAT3 stabilizes KLF4 protein in ES cells and is involved in recruiting KLF4 to the RNAPII compartment.

**Figure 5:**
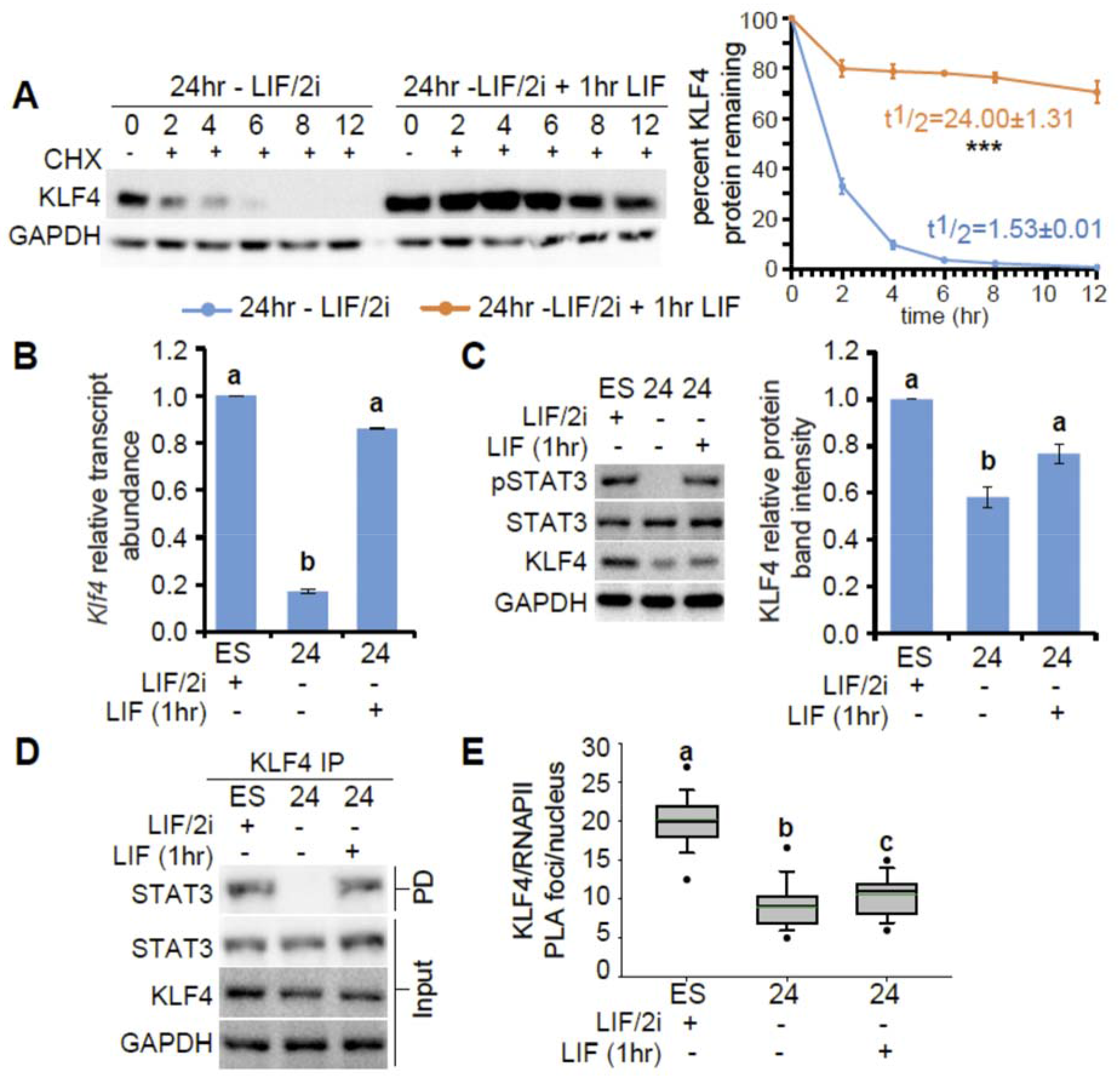
Phosphorylated STAT3 stabilizes KLF4 protein in nuclear complexes. A) Immunoblots for KLF4 and GAPDH in ES cells 24hr after LIF/2i removal and in cells 24hr after LIF/2i removal followed by a 1hr pre-treatment with LIF, sampled at 0, 2, 4, 6, 8 and 12 hr after CHX treatment. GAPDH levels were used as a control and displayed the expected half-life (t_½_ >30hr). On the right percent remaining KLF4 was calculated from the intensity of CHX treatment immunoblots, measured in three biological replicates. Half-life was calculated for each time series replicate by best fit to exponential decay. Error bars represent standard deviation. Statistical differences between protein half-life determined by two-tailed t-test (P < 0.001) are indicated as ***. B) Transcript levels for *Klf4* were quantified relative to *Gapdh* and ES levels, in three biological replicates of ES cells cultured with LIF/2i, 24hr after LIF/2i removal and in cells 24hr after LIF/2i removal followed by a 1hr treatment with LIF. Error bars represent standard deviation. Statistical differences were determined by one-way ANOVA (P < 0.05) and displayed by different letters. C) Immunoblots for pSTAT3 (Tyr705) and total STAT3 in ES cells cultured with LIF/2i, 24hr after LIF/2i removal and in cells 24hr after LIF/2i removal followed by a 1hr treatment with LIF. GAPDH levels indicate sample loading. On the right KLF4 protein relative to GAPDH and ES levels, in three biological replicates of ES cells cultured with LIF/2i, 24hr after LIF/2i removal and in cells 24hr after LIF/2i removal followed by a 1hr treatment with LIF. Error bars represent standard deviation. Statistical differences determined with one-way ANOVA (P < 0.05) are displayed by different letters. D) KLF4 immunoprecipitation (IP) in ES cells cultured with LIF/2i, 24hr after LIF/2i removal and in cells 24hr after LIF/2i removal followed by a 1hr treatment with LIF, probed with anti-STAT3 (PD, pull down). GAPDH levels indicate sample loading of the input. E) Box-and-whisker plots display the number of KLF4/RNAPII PLA foci per nucleus for ES cells cultured with LIF/2i, 24hr after LIF/2i removal and in cells 24hr after LIF/2i removal followed by a 1hr treatment with LIF. Boxes indicate interquartile range of intensity values and whiskers indicate the 10th and 90th percentiles; outliers are shown as black dots. Images were collected from at least three biological replicates and ≥100 nuclei were quantified for each sample. Statistical differences determined with one way ANOVA (P < 0.05) are indicated by different letters.

### SOX2 and NANOG regulate KLF4 post-translationally by stabilizing KLF4 protein in nuclear complexes

As association with pSTAT3 increased the stability of KLF4 protein after a 1hr treatment with LIF, we next investigated whether additional pluripotency transcription factors, SOX2 and NANOG, had a role in controlling KLF4 stability. Cells with compromised SOX2 protein levels due to homozygous deletion of the SCR express reduced levels of KLF4 protein but not RNA (Figure 1) which could be due to reduced KLF4 protein stability in these cells. The ΔSCR^129/Cast^ cells are female F1 ES cells which are partially differentiated; they have inactivated one X chromosome and display upregulation of genes normally expressed in trophoblast stem cells, determined by RNA-seq (Zhou et al., 2014). In addition, we showed that the levels of *Oct4, Nanog, Klf2* and *Klf5* transcripts as well as protein were not affected by deletion of the SCR (Figure S1).

Evaulating KLF4 protein stability in ΔSCR^129/Cast^ cells revealed that KLF4 is unstable with a t_½_ <2hrs and this reduced stability is unaffected by removal of LIF/2i (Figure 6A). To evaluate the role of SOX2 in stabilizing KLF4 protein, ΔSCR^129/Cast^ cells were transfected with Sox2-t2A-GFP. Transfection restored *Sox2* mRNA and protein to wild type levels and significantly increased KLF4 protein but not transcript levels, indicating regulation of KLF4 occurs post-transcriptionally by SOX2 (Figure 6B/C). In addition, the stability of KLF4 protein increased significantly from t_½_=2hr to t_½_=17hr after SOX2 protein levels were restored (Figure 6D). To investigate whether SOX2 could stabilize KLF4 through protein-protein interaction, KLF4 was immunoprecipitated to investigate its interaction with SOX2. Indeed KLF4 and SOX2 do interact and this interaction is restored in Sox2-t2A-GFP transfected cells (Figure 6E). Similarly, increased nuclear association between KLF4 and SOX2 was observed by PLA in Sox2-t2A-GFP transfected cells (Figure 6F). Although, the number of PLA foci/nucleus for KLF4 with RNAPII-S5P was unchanged when SOX2 protein levels were restored, the signal intensity per focus increased significantly, suggesting SOX2 protein is involved in stabilizing KLF4 protein in nuclear transcriptional complexes (Figure 6G).

**Figure 6:**
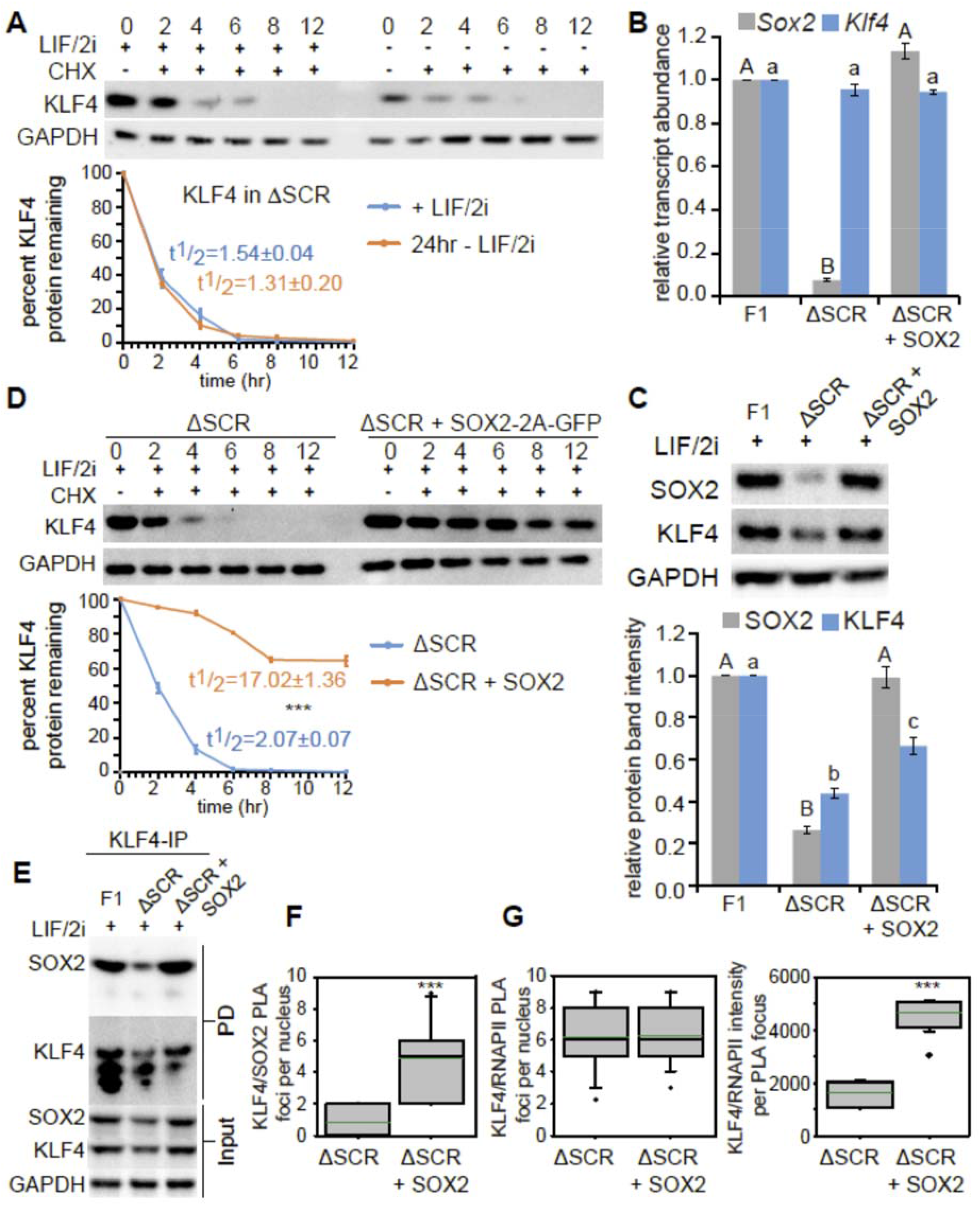
SOX2 stabilizes KLF4 protein in nuclear complexes. A) Immunoblots for KLF4 in the ΔSCR^129/Cast^ clone (ΔSCR), cultured with LIF/2i and 24hr after removal of LIF/2i, sampled at 0, 2, 4, 6, 8 and 12hr after CHX treatment. GAPDH levels were used as a control and displayed the expected half-life (t_½_ >30hr). The percent remaining KLF4 protein was calculated from the intensity of CHX treatment immunoblots, measured in three biological replicates. KLF4 protein half-life in the presence or absence of LIF/2i was calculated for each time series replicate by best fit to exponential decay. Error bars represent standard deviation of three biological replicates. Statistical differences determined by two-tailed t-test are indicated by *** (P < 0.001). B) *Sox2* and *Klf4* transcripts of were quantified relative to *Gapdh* and F1, in three biological replicates of F1, ΔSCR^129/Cast^ and SOX2-2A-GFP transfected ΔSCR^129/Cast^. Error bars represent standard deviation. Statistical differences for each transcript were determined by one way ANOVA (P < 0.05) and displayed as upper case letters for *Sox2* and lower case letters for *Klf4*. C) Immunoblots for SOX2 and KLF4 in F1, ΔSCR^129/Cast^ and SOX2-2A-GFP transfected ΔSCR^129/Cast^ cells in LIF/2i. GAPDH levels indicate sample loading. Quantification of KLF4 and SOX2 relative to GAPDH and F1, in three biological replicates. Error bars represent standard deviation. Statistical differences for each protein were determined by one way ANOVA (P < 0.05) and displayed as upper case letters for SOX2 and lower case letters for KLF4. D) Immunoblot for KLF4 and GAPDH in untransfected and SOX2-2A-GFP transfected ΔSCR^129/Cast^ cells in LIF/2i sampled at 0, 2, 4, 6, 8 and 12hr after CHX treatment. GAPDH levels were used as a control and displayed the expected half-life (t_½_ >30hr). Percent remaining KLF4 protein at 0, 2, 4, 6, 8, and 12hr was calculated from the intensity of CHX treatment immunoblots, measured in three biological replicates. Half-life was calculated for each time series replicate by best fit to exponential decay. Error bars represent standard deviation. Statistical differences determined by two-tailed t-test (P < 0.001) are indicated as ***. E) KLF4 immunoprecipitation (IP) from F1 ES, untransfected and SOX2-2A-GFP transfected ΔSCR^129/Cast^ cells probed with anti-SOX2 and anti-KLF4 (PD, pull down). GAPDH levels indicate loading of the input. F-G) Box-and-whisker plots boxes indicate interquartile range of intensity values and whiskers indicate the 10th and 90th percentiles; outliers are shown as black dots. Images were collected from at least three biological replicates and ≥100 nuclei were quantified for each sample. Statistical differences determined by t test (P < 0.001) are indicated as ***. In F the number of KLF4/SOX2 PLA foci per nucleus are shown. In G the number of KLF4/RNAPII PLA foci per nucleus are shown (left) and the total intensity value of each PLA focus is shown (right).

As the levels of NANOG protein in the nucleus are drastically reduced by 24hr of differentiation, whereas SOX2 levels remain high at this time (Dhaliwal et al., 2018), we hypothesized that loss of NANOG protein, in addition to the loss of pSTAT3, could be involved in reduced KLF4 stability after 24hr of differentiation. To test this hypothesis, cells differentiated for 24hr were transfected with Nanog-t2a-GFP to restore NANOG levels. After an additional 24hr in differentiation media, KLF4 protein levels were restored in Nanog-t2a-GFP transfected cells without any change in *Klf4* transcript levels, indicating a post-transcriptional regulatory mechanism (Figure 7A/B). We also monitored the expression of *Oct4, Sox2, Klf2* and *Klf5* RNA and protein and found that they were not altered during this differentiation timeframe or by Nanog-t2a-GFP transfection (Figure S8). Similar to what was observed for SOX2, NANOG expression in differentiated cells restores KLF4 protein stability from t_½_ = 1.5hr to t_½_ =25hr (Figure 7C). To investigate whether NANOG could stabilize KLF4 through protein-protein interaction, KLF4 was immunoprecipitated to investigate interaction with NANOG; indeed KLF4 and NANOG do interact in differentiated cells transfected with Nanog-t2A-GFP (Figure 7D). Restoring NANOG levels in differentiated cells not only increased the interaction between KLF4 and NANOG but also restored the interaction of KLF4 with RNAPII-S5P that is normally lost by 48hr of differentiation, as revealed by PLA (Figure 7E). Together these data reveal that NANOG has a role in both stabilizing KLF4 protein and recruiting KLF4 to RNAPII compartments in the nucleus.

**Figure 7:**
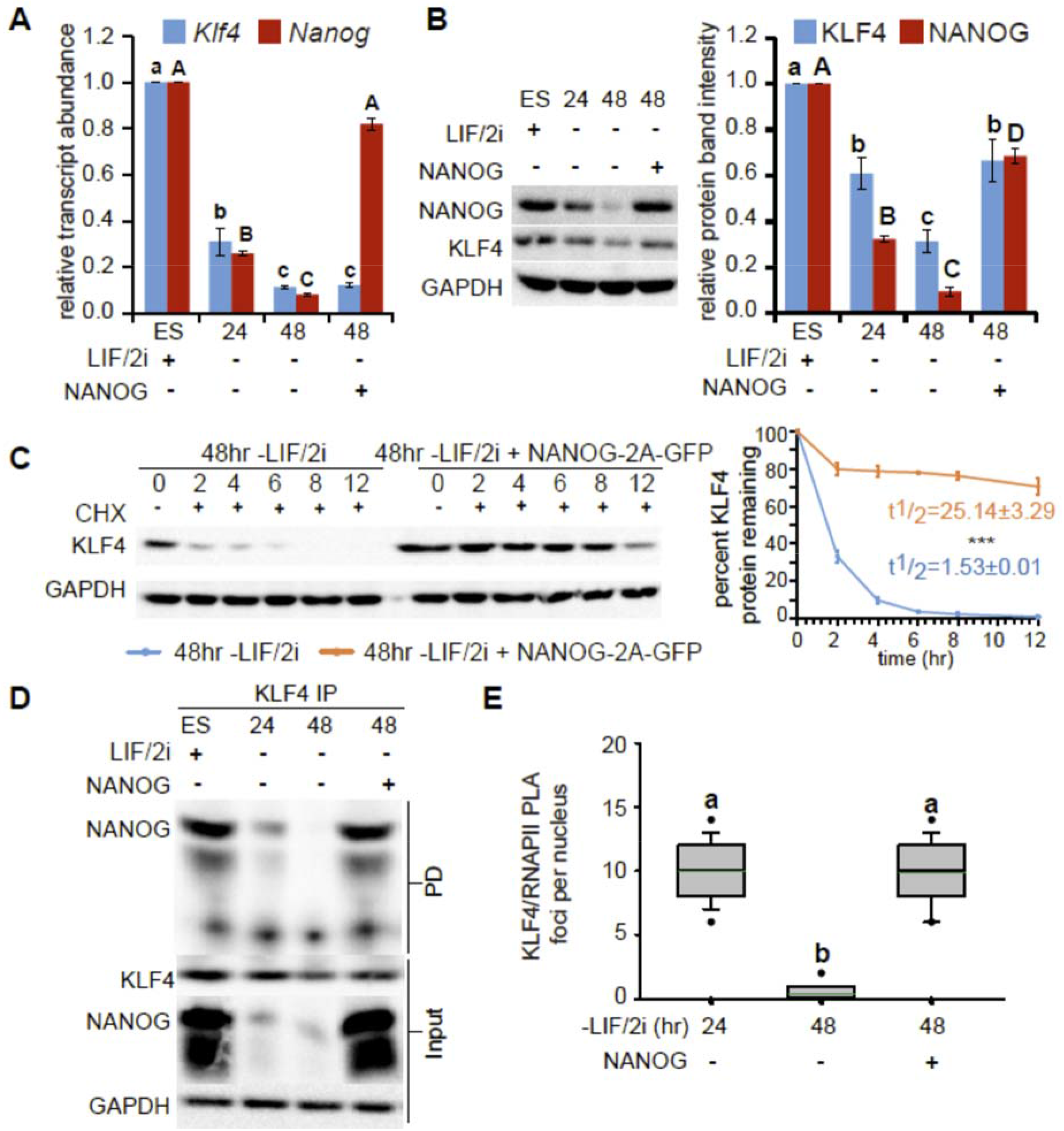
NANOG stabilizes KLF4 protein in nuclear complexes. A) Transcript levels of *Klf4* and *Nanog* were quantified relative to *Gapdh* and undifferentiated ES cells in three biological replicates of ES cells cultured with LIF/2i (ES), 24 and 48hr after removal of LIF/2i, and in ES cells 48hr after removal of LIF/2i where NANOG-2A-GFP was transfected 24hr after removal of LIF/2i. Error bars represent standard deviation. Statistical differences for each transcript were determined by one way ANOVA (p<0.05) and displayed as lower case letters for *Klf4* and upper case letters for *Nanog*. B) Immunoblots for KLF4 and NANOG in ES cells cultured with LIF/2i, 24 and 48hr after removal of LIF/2i, and in ES cells 48hr after removal of LIF/2i where NANOG-2A-GFP was transfected 24hr after removal of LIF/2i. GAPDH levels indicate sample loading. On the right quantification of KLF4 and NANOG relative to GAPDH for three biological replicates. Error bars represent standard deviation. Statistical differences for each protein were determined by one way ANOVA (p<0.05) and displayed as lower case letters for KLF4 and upper case letters for NANOG. C) Immunoblot for KLF4 and GAPDH in ES cells 48hr after removal of LIF/2i, and in ES cells 48hr after removal of LIF/2i where NANOG-2A-GFP was transfected 24hr after removal of LIF/2i, sampled at 0, 2, 4, 6, 8 and 12 hr after CHX treatment. GAPDH levels were used as a control and displayed the expected half-life (t_½_ >30hr). On the right the percent remaining KLF4 was calculated from the intensity of CHX treatment immunoblots, measured in three biological replicates. Half-life was calculated for each time series replicate by best fit to exponential decay. Error bars represent standard deviation. Statistical differences between protein half-life determined by two-tailed t-test (P < 0.001) are indicated as ***. D) KLF4 immunoprecipitation (IP) from ES cells cultured with LIF/2i, 24 and 48hr after removal of LIF/2i, and in ES cells 48hr after removal of LIF/2i where NANOG-2A-GFP was transfected 24hr after removal of LIF/2i, probed with anti-NANOG (PD, pull down). GAPDH levels indicate loading of the input. E) Box-and-whisker plots indicate the number PLA foci per nucleus for the interaction between KLF4 and RNAPII. Boxes indicate interquartile range of intensity values and whiskers indicate the 10th and 90th percentiles; outliers are shown as black dots. Images were collected from at least three biological replicates and ≥100 nuclei were quantified for each sample. Statistical differences determined by one-way ANOVA (P < 0.05) are indicated by different letters.

To investigate the role of pluripotency associated transcription factors as stabilizing agents of KLF4 protein in another cellular context we transfected HEK293 cells (which do not express these pluripotency regulators) with KLF4-GFP. KLF4-GFP displayed low protein stability in HEK293 cells with a t_½_ ~1hr (Figure S8). Co-expression of SOX2 increased KLF4 protein stability; however, SOX2 was not able to increase the stability of KLF4ΔZNF-GFP, indicating the DNA binding zinc finger domains are important for stabilization of KLF4 by SOX2. This implicates protein-protein interaction between SOX2 and KLF4 as a key mechanism as SOX2 has been shown to associate with KLF4 through the KLF4 zinc finger domains (Wei et al., 2009). By contrast, both WT KLF4-GFP and KLF4ΔZNF-GFP were stabilized to a similar extent by a complex of SOX2, NANOG and constitutively active STAT3 (CA-STAT3) and removal of SOX2 from this complex did not alter KLF4 stability (Figure S8). Addition of KLF2 to the full complex or co-transfection of only KLF2 with KLF4-GFP did not further stabilize KLF4 protein indicating there is no direct role for KLF2 in stabilizing KLF4 in this context.

Together these data indicate that nuclear anchoring, through interaction with RNAPII complexes, pluripotency transcription factors, and association with DNA, maintains the stability of KLF4 protein in undifferentiated naïve pluripotent ES cells.

## Discussion

The naïve pluripotent state of ES cells is regulated by pluripotency associated transcription factors, which act in a complex interconnected network by binding their own regulatory elements, as well as the regulatory elements of other genes throughout the genome to regulate transcription. *Klf4* is regulated at a transcriptional level by downstream enhancers which are required to maintain *Klf4* transcript and protein levels in mouse ES cells maintained in LIF/serum (Xie et al., 2017). By contrast, we found that for ES cells maintained in LIF/2i the same enhancers had little role in maintaining KLF4 protein levels as the high stability of the KLF4 protein in LIF/2i conditions buffered against the greatly reduced levels of *Klf4* transcription in the absence of the enhancers. Removal of MEK inhibition greatly disrupted KLF4 stability, accounting for the dramatic differences between the requirements for *Klf4* enhancers in LIF/serum compared to LIF/2i conditions. Furthermore, we found that KLF4 protein is stabilized both by domains within the protein that anchor KLF4 in the nucleus and by interaction with RNAPII, SOX2, NANOG and pSTAT3. These findings detail a new way in which the function of the core pluripotency regulatory circuitry is integrated at a post-translational level by controlling KLF4 protein stability. Furthermore, as we found that both SOX2 and NANOG regulated KLF4 only by post-translational mechanisms our findings contradict the accepted regulatory circuitry which states that expression of these transcription factors is integrated by transcriptional mechanisms.

Nuclear localization and retention of KLF4 plays a key role in its regulation. KLF4 requires an intact NLS for nuclear import as NLS mutant KLF4 protein is cytoplasmic (Dhaliwal et al., 2018). KLF4 nuclear localization increases protein stability as the NLS mutant displayed greatly reduced stability compared to the wild type protein. Mutations which disrupted KLF4 nuclear anchoring also disrupted KLF4 protein stability in LIF/2i conditions. Deletion of the KLF4 DNA binding zinc finger domains or mutation of the K275 sumoylation site, involved in transactivation, caused a partial disruption to KLF4 nuclear anchoring. An important consequence of these mutations is that the stability of the protein was completely abolished, suggesting that even a small disruption to nuclear anchoring can greatly affect KLF4 protein stability. By contrast, in COS-1 cells expression of a KLF4 K275 mutant did not affect protein stability further indicating that regulation of KLF4 protein stability is context dependent (Du et al., 2010). The transactivation function of KLF4 has been shown to be regulated in a SUMO1-dependent manner; mutation of the K275 sumoylation site in mouse KLF4 impairs the ability of KLF4 to transactivate target promoters by disrupting the SUMO-interacting motif in KLF4 (Du et al., 2010). We found that this mutation also reduced recruitment of KLF4 to RNAPII rich nuclear compartments which would explain reduced transactivation of the K275 mutant protein. Restoring nuclear anchoring in this transactivation mutant did not restore interaction with RNAPII indicating that K275 is indeed required for interaction with RNAPII and KLF4 function.

ERK activation-induced KLF4 nuclear export is a critical first step in pluripotency exit which relies on KLF4 phosphorylation at S132 and intact NES allowing for interaction with XPO1 (Dhaliwal et al., 2018). We determined that mono-ubiquitination of KLF4 at K249 is also required for KLF4 nuclear export and pluripotency exit although this event occurs later after 24hr of differentiation. Three regions, S132, K249 and the KLF4 NES, are required for the disruption to KLF4 stability that occurs as ES cells exit the pluripotent state and preventing KLF4 phosphorylation, ubiquitination or interaction with XPO1 blocks pluripotency exit which normally occurs after removal of LIF/2i. By contrast, even in the presence of LIF/2i, KLF4 stability can be disrupted by interfering with KLF4 function by deletion of the DNA binding zinc finger domains or mutation of the K275 sumoylation site, involved in transactivation. The compound mutants (S132AΔZNF, S132AK275R), however, maintain KLF4 stability but not KLF4 regulatory activity or interaction with RNAPII and therefore do not block ES cell differentiation.

Naïve pluripotent ES cells cultured in LIF/2i are thought to maintain a more balanced and stable state than ES cells maintained in LIF/serum, through higher and more uniform expression of the major pluripotency transcription factors in the cell population (Sim et al., 2017; Tosolini and Jouneau, 2016). Our results indicate that for KLF4 protein this higher and more uniform expression in LIF/2i is achieved through post-translational protein stabilization. This in turn could have knock-on effects leading to higher transcription of additional pluripotency associated transcription factors. Protein interactome studies determined that the core pluripotency transcription factors bind each other and form nuclear complexes in order to maintain the pluripotent state (Gao et al., 2013; Gao et al., 2012; Morey et al., 2015). KLF4 protein stability is maintained in ES cells through interaction of KLF4 with pSTAT3, NANOG, and SOX2 in RNAPII rich nuclear complexes. Co-immunoprecipitation revealed that these interactions are maintained in the absence of both DNA and RNA as these were removed by nuclease treatment during co-immunoprecipitation, suggesting they are protein-protein interactions in nature. Even though KLF4-GFP co-expressed with SOX2, NANOG and CA-STAT3 in HEK293 cells displayed higher protein stability than KLF4-GFP expressed alone, the maximal HEK293 stability observed for KLF4-GFP (t_½_ 3hr) remained lower than what was observed in undifferentiated ES cells (t_½_ 24hr) suggesting additional, unknown factors are involved in context dependent stabilization of the KLF4 protein in pluripotent cells. KLF4 is exceptional in that the stability of other pluripotency factors, or the other expressed *Klf* family members, are not affected in the same way by these complexes as their stability does not change during pluripotency exit. Although KLF4 is not required during mouse development (Segre et al., 1999), its downregulation through a disruption in protein satbility is likely important during development as we found that preventing protein destabilization blocks pluripotency exit and differentiation. Furthermore, the mutations to the KLF4 protein that increase KLF4 stability, without affecting KLF4 function as a transcription factor, could be deployed in a reprogramming context to improve the efficiency of reprogramming protocols by prolonging KLF4 protein function in reprogramming cells.

In addition to the role of KLF4 protein in pluripotency maintenance and reprogramming, KLF4 protein modulation is involved in both the oncogenic and tumor suppressor roles of KLF4 in adult tissues. Increased stability of the KLF4 protein, mediated by the ubiquitin-proteosomal pathway, is linked to an oncogenic role in breast cancer (Hu et al., 2012) and decreased stability is linked to its tumor suppressor role in colorectal cancer (Gamper et al., 2012). KLF4 stability modulation by interaction with the pluripotency transcription factors may also be important in tumorigenesis as co-expression of these factors is associated with a variety of cancers such as breast cancer, squamous cell carcinoma, and gastrointestinal cancer progression (Almozyan et al., 2017; Gwak et al., 2017; Hu et al., 2012; Lu et al., 2014; Piva et al., 2014; Soheili et al., 2017).

## Materials and Methods

### Embryonic stem Cell culture

The mouse ES cell line E14TG2a (E14) was obtained from the ATCC (CRL-1821). F1 (*M. musculus*^129^ × *M. castaneus*) ES cells were obtained from Barbara Panning (Mlynarczyk-Evans et al., 2006). All cells were maintained in feeder free conditions on 0.1% gelatin in DMEM supplemented with 15% (v/v) fetal bovine serum (FBS), 0.1mM non-essential amino acids, 1mM sodium pyruvate, 2mM GlutaMAX, 0.1mM 2-mercaptoethanol, 1,000U/mL LIF, 3μM CHIR99021 (GSK3β inhibitor, Biovision) and 1μM PD0325901 (MEK inhibitor, Invivogen); referred to as LIF/2i medium. 18 hr prior to differentiation the cells were seeded at 10,000 cells/cm^2^. The differentiation medium contained the same components with the exception of LIF and the two inhibitors.

For protein half-life analysis cells were treated with 10μg/ml cyclohexamide (Sigma Aldrich) for 2, 4 6, 8 and 12hr, collected as cell pellets, and lysed in RIPA buffer containing protease inhibitor complete EDTA free (Roche) and phosphatase inhibitor cocktail (Millipore) to generate cell lysates for further analysis by western blotting. Treatment with 10μM MG132 proteasome inhibitor (Sigma Aldrich) was used to indicate the role of proteosomal degradation in KLF4 function.

### Cellular fractionation, co-immunoprecipitation and western blotting

Nuclear and cytoplasmic fractions were generated according to the protocol described in (Dhaliwal and Mitchell, 2016). Protein was extracted from cell fractions using RIPA buffer containing protease inhibitor complete EDTA free (Roche) and phosphatase inhibitor cocktail (Millipore) and quantified using bicinchoninic acid (Thermo Fisher Scientific). Protein samples were analyzed by SDS-PAGE (Bis-Tris, 5% stacking, 10% resolving). Blots were probed with primary antibodies followed by horseradish peroxidase-conjugated secondary antibodies (Table S1). Blots were quantified by relative intensity using background correction from adjacent regions. At least three biological replicates were analyzed for each experiment.

For co-immunoprecipitation of protein, fractions or total cell lysates in RIPA were treated with Benzonase nuclease (Sigma-Aldrich) to remove RNA and DNA and then incubated overnight with the appropriate antibody and then incubated overnight with a 50:50 mixture of protein A and protein G Dynabeads (Thermo Fisher Scientific). Beads were washed three times with nondenaturing lysis buffer (20 mM Tris-HCl, 137 mM NaCl, 10% glycerol, 1% NP-40, 2 mM EDTA, 1 mM PMSF, and proteinase inhibitors), twice with PBS, eluted in SDS-PAGE loading buffer, and analyzed by SDS-PAGE. Antibodies used are listed in table S1.

### Proximity Ligation Amplification (PLA)

PLA was conducted using Duolink (Sigma-Aldrich) following the manufacturer’s instructions. Images were collected using a Leica TCS SP8 and a 63× magnification objective lens. The number of PLA foci per nucleus was quantified using Imaris 7.1 by manual 3D masking of nuclei in ESC colonies defined by the DAPI signal. All PLA experiments were carried out on at least three biological replicate samples.

### CRISPR/Cas9 mediated deletion

Cas9-mediated deletions were carried out as described (Moorthy and Mitchell, 2016; Zhou et al., 2014). Cas9 targeting guides flanking *Klf4* enhancer regions were selected (Table S2). Only gRNAs predicted to have no off-target binding in the F1 mouse genome were chosen. Guide RNA plasmids were assembled in the gRNA vector (Addgene, ID#41824) using the protocol described by (Mali et al., 2013). The sequence of the resulting guide plasmid was confirmed by sequencing. F1 ES cells were transfected with 5 μg each of 5′ gRNA, 3′ gRNA, and pCas9_GFP (Addgene, ID#44719)(Ding et al., 2013) plasmids using the Neon Transfection System (Life Technologies). Forty-eight hours post-transfection, GFP-positive cells were collected and sorted on a BD FACSAria. Ten to twenty thousand GFP positive cells were seeded on 10-cm gelatinized culture plates and grown for 5–6 d until large individual colonies formed. Colonies were picked and propagated for genotyping and gene expression analysis. All deletions were confirmed by sequence analysis using primers 5′ and 3′ from the gRNA target sites; SNPs within the amplified product confirmed the genotype of the deleted allele. Transcription factor binding at the *Klf4* locus was determined by evaluating data obtained from CODEX (Sanchez-Castillo et al., 2014). ChIP-seq data for KLF5 was obtained from Aksoy et al. 2014 (Aksoy et al., 2014).

### Real-Time qPCR

Total RNA was purified as per manufacturer’s protocol using RNAeasy (Qiagen). Following a DNaseI (turbo DNaseI from Invitrogen) digestion to remove DNA, total RNA was reverse transcribed with random primers using the High Capacity cDNA Synthesis Kit (Applied biosystems). Gene expression was monitored by qPCR using genomic DNA to generate standard curves. Gapdh expression was used to normalize expression values. At least three biological replicates were analyzed for each experiment. Primers used are listed in Table S3. All samples were confirmed not to have DNA contamination by generating a reverse transcriptase negative sample and monitoring *Gapdh* amplification. Allele specific primers were determined to amplify only the specified allele by testing amplification on C57BL/6 (same genotype as 129 at the target sequence) and castaneus genomic DNA.

### Expression of KLF4 Mutants

A mouse KLF4-GFP vector (RG206691) obtained from Origene and KLF4(S132A)-GFP mutant published in Dhaliwal et al. 2018 were subjected to site-directed mutagenesis (SDM, QuikChange Lightning, Agilent Technologies) to introduce additional mutations. Primers for SDM are indicated in Table 4. A MluI/AleI restriction digestion deleted zinc fingers from the *Klf4* sequence cloned in the pUC 19 vector. *Klf4* in pUC19 without Zinc fingers (3881bp) was ligated by blunt end ligation. After sequence confirmation, the zinc finger deleted *Klf4* fragment was inserted into a Kpn1/Not1 digested KLF4-GFP vector. Sequence-confirmed plasmids were transfected by electroporation into E14 ES cells and selected with 400 μg/mL G418. The cells were sorted by fluorescence-activated cell sorting, and individual clones selected and maintained in 50 μg/mL G418 to obtain KLF4-GFP-positive clones.

### Transient transfections

Sox2-t2A-GFP was generated by amplifying the human *SOX2* sequence from a donor construct (HsCD00079917, Harvard Institute of Proteomics)(Zuo et al., 2007) using NheI and XbaI overhang primers and Phusion High-Fidelity Polymerase (NEB). A t2a-GFP backbone was subcloned from pLV hU6-sgRNA hUbC-dCas9-KRAB-t2a-GFP (Addgene plasmid #71237, a gift from Charles Gersbach)(Thakore et al., 2015) and inserted into hCas9 (Addgene plasmid #41815, a gift from George Church)(Mali et al., 2013) using AgeI/XbaI digestion and T4 DNA ligase (Thermo Fisher Scientific). Next, dCas9-KRAB was excised out using NheI/XbaI to create a lineralized CMV-t2a-GFP construct. The *SOX2* PCR product was purified and inserted into CMV-t2a-GFP using the In-Fusion HD cloning kit (Takara Bio USA) and transformed into Stellar Competent Cells (Takara Bio USA). To produce Sox2-t2A-mCherry, Sox2-t2A-GFP was then cut with KpnI and AgeI to remove GFP and replace with PCR-amplified mCherry from pAAVS1-NDi-CRISPRi (Addgene plasmid #73497, a gift from Bruce Conklin)(Mandegar et al., 2016) with KpnI and AgeI overhang primers. Sox2-t2A-mCherry was digested with XbaI and NheI to remove *Sox2* and replace with PCR products from *Nanog* (Addgene plasmid # 59994, a gift from Rudolf Jaenisch)(Faddah et al., 2013), *Klf2* (Addgene plasmid #66655, a gift from Barak Cohen) and CA-*Stat3* (Addgene plasmid #8722, a gift from Jim Darnell)(Bromberg et al., 1999). Bacterial colonies were PCR-screened, and positive inserts were sequence confirmed. All plasmis were purified with an Endotoxin-free Plasmid Midiprep Kit (Geneaid™ Midi Plasmid Kit Endotoxin Free).

The SOX2 compromised SCR^129/Cast^ deleted cells were transfected with Sox2-t2A-GFP and 24hr differentiated ES cells were transfected with Nanog-t2A-GFP using neon electroporation transfection system as per manufacturer’s instructions (Thermo Fisher Scientific). Nanog-2A-GFP was a gift from Rudolf Jaenisch (Addgene plasmid # 59994)(Faddah et al., 2013).

HEK293 (Flp-In™-293 Cell Line, Thermo Fisher Scientific) were grown in DMEM supplemented with 10% FBS, 1x PenStrep, 1x Glutamax, 1x Non-essential amino acids, 1x Sodium Pyruvate. Cells were split into 6-well plates and seeded at 30,000 cells/cm^2^ 18hr before transfection. KLF4-GFP was transfected alone or with Klf2-t2A-mCherry, Sox2-t2A-mCherry, CA-Stat3-t2A-mCherry or Nanog-t2A-mCherry in different combinations at 2μg per well. Lipofectamine 3000 at 0.3 μl per μl of Opti-MEM (Thermo Fisher Scientific) was mixed with plasmids at 1.5μl/μg of plasmid. Transfected cells were treated with 10 μg/ml CHX (Sigma Aldrich) and sampled after 0, 2,4,6,8 and 12 hrs.

## Acknowledgements

This work was supported by the Canadian Institutes of Health Research (FRN 153186), the Canada Foundation for Innovation, and the Ontario Ministry of Research and Innovation (operating and infrastructure grants held by J.A.M.). Studentship funding was provided by Ontario Graduate Scholarships held by N.K.D. The authors declare no competing interests.

## Author contributions

J.A.M. and N.K.D. conceived and designed the experiments. N.K.D. performed all experiments with the following assistance: L.E.A. constructed several of the vectors used in this study and analyzed KLF5 ChIP-seq data. N.K.D. and J.A.M. analyzed the data and wrote the manuscript, which was approved by all co-authors.

## REFERENCES

Aksoy, I., Giudice, V., Delahaye, E., Wianny, F., Aubry, M., Mure, M., Chen, J., Jauch, R., Bogu, G.K., Nolden, T., et al. (2014). Klf4 and Klf5 differentially inhibit mesoderm and endoderm differentiation in embryonic stem cells. Nat Commun 5, 3719.

Almozyan, S., Colak, D., Mansour, F., Alaiya, A., Al-Harazi, O., Qattan, A., Al-Mohanna, F., Al-Alwan, M., and Ghebeh, H. (2017). PD-L1 promotes OCT4 and Nanog expression in breast cancer stem cells by sustaining PI3K/AKT pathway activation. Int J Cancer 141, 1402–1412.

Blinka, S., Reimer, M.H., Jr., Pulakanti, K., and Rao, S. (2016). Super-Enhancers at the Nanog Locus Differentially Regulate Neighboring Pluripotency-Associated Genes. Cell Rep 17, 19–28.

Bromberg, J.F., Wrzeszczynska, M.H., Devgan, G., Zhao, Y., Pestell, R.G., Albanese, C., and Darnell, J.E., Jr. (1999). Stat3 as an oncogene. Cell 98, 295–303.

Carter, D., Chakalova, L., Osborne, C.S., Dai, Y.F., and Fraser, P. (2002). Long-range chromatin regulatory interactions in vivo. Nat Genet 32, 623–626.

Chen, C.Y., Morris, Q., and Mitchell, J.A. (2012). Enhancer identification in mouse embryonic stem cells using integrative modeling of chromatin and genomic features. BMC Genomics 13, 152.

Chen, X., Xu, H., Yuan, P., Fang, F., Huss, M., Vega, V.B., Wong, E., Orlov, Y.L., Zhang, W., Jiang, J., et al. (2008). Integration of external signaling pathways with the core transcriptional network in embryonic stem cells. Cell 133, 1106–1117.

Dhaliwal, N.K., Miri, K., Davidson, S., Tamim El Jarkass, H., and Mitchell, J.A. (2018). KLF4 Nuclear Export Requires ERK Activation and Initiates Exit from Naive Pluripotency. Stem Cell Reports 10, 1308–1323.

Dhaliwal, N.K., and Mitchell, J.A. (2016). Nuclear RNA Isolation and Sequencing. Methods Mol Biol 1402, 63–71.

Ding, Q., Regan, S.N., Xia, Y., Oostrom, L.A., Cowan, C.A., and Musunuru, K. (2013). Enhanced efficiency of human pluripotent stem cell genome editing through replacing TALENs with CRISPRs. Cell Stem Cell 12, 393–394.

Du, J.X., McConnell, B.B., and Yang, V.W. (2010). A small ubiquitin-related modifier-interacting motif functions as the transcriptional activation domain of Kruppel-like factor 4. J Biol Chem 285, 28298–28308.

Faddah, D.A., Wang, H., Cheng, A.W., Katz, Y., Buganim, Y., and Jaenisch, R. (2013). Single-cell analysis reveals that expression of nanog is biallelic and equally variable as that of other pluripotency factors in mouse ESCs. Cell Stem Cell 13, 23–29.

Gamper, A.M., Qiao, X., Kim, J., Zhang, L., DeSimone, M.C., Rathmell, W.K., and Wan, Y. (2012). Regulation of KLF4 turnover reveals an unexpected tissue-specific role of pVHL in tumorigenesis. Mol Cell 45, 233–243.

Gao, F., Wei, Z., An, W., Wang, K., and Lu, W. (2013). The interactomes of POU5F1 and SOX2 enhancers in human embryonic stem cells. Sci Rep 3, 1588.

Gao, Z., Cox, J.L., Gilmore, J.M., Ormsbee, B.D., Mallanna, S.K., Washburn, M.P., and Rizzino, A. (2012). Determination of protein interactome of transcription factor Sox2 in embryonic stem cells engineered for inducible expression of four reprogramming factors. J Biol Chem 287, 11384–11397.

Gwak, J.M., Kim, M., Kim, H.J., Jang, M.H., and Park, S.Y. (2017). Expression of embryonal stem cell transcription factors in breast cancer: Oct4 as an indicator for poor clinical outcome and tamoxifen resistance. Oncotarget 8, 36305–36318.

Hall, J., Guo, G., Wray, J., Eyres, I., Nichols, J., Grotewold, L., Morfopoulou, S., Humphreys, P., Mansfield, W., Walker, R., et al. (2009). Oct4 and LIF/Stat3 additively induce Kruppel factors to sustain embryonic stem cell self-renewal. Cell Stem Cell 5, 597–609.

Hirai, H., Karian, P., and Kikyo, N. (2011). Regulation of embryonic stem cell self-renewal and pluripotency by leukaemia inhibitory factor. Biochem J 438, 11–23.

Hochstrasser, M., and Varshavsky, A. (1990). In vivo degradation of a transcriptional regulator: the yeast alpha 2 repressor. Cell 61, 697–708.

Hu, D., Zhou, Z., Davidson, N.E., Huang, Y., and Wan, Y. (2012). Novel insight into KLF4 proteolytic regulation in estrogen receptor signaling and breast carcinogenesis. J Biol Chem 287, 13584–13597.

Jovanovic, M., Rooney, M.S., Mertins, P., Przybylski, D., Chevrier, N., Satija, R., Rodriguez, E.H., Fields, A.P., Schwartz, S., Raychowdhury, R., et al. (2015). Immunogenetics. Dynamic profiling of the protein life cycle in response to pathogens. Science 347, 1259038.

Kesmir, C., Nussbaum, A.K., Schild, H., Detours, V., and Brunak, S. (2002). Prediction of proteasome cleavage motifs by neural networks. Protein Eng 15, 287–296.

Kim, M.O., Kim, S.H., Cho, Y.Y., Nadas, J., Jeong, C.H., Yao, K., Kim, D.J., Yu, D.H., Keum, Y.S., Lee, K.Y., et al. (2012). ERK1 and ERK2 regulate embryonic stem cell self-renewal through phosphorylation of Klf4. Nat Struct Mol Biol 19, 283–290.

Kim, S.H., Kim, M.O., Cho, Y.Y., Yao, K., Kim, D.J., Jeong, C.H., Yu, D.H., Bae, K.B., Cho, E.J., Jung, S.K., et al. (2014). ERK1 phosphorylates Nanog to regulate protein stability and stem cell self-renewal. Stem Cell Res 13, 1–11.

Kristensen, A.R., Gsponer, J., and Foster, L.J. (2013). Protein synthesis rate is the predominant regulator of protein expression during differentiation. Mol Syst Biol 9, 689.

Li, Y., Rivera, C.M., Ishii, H., Jin, F., Selvaraj, S., Lee, A.Y., Dixon, J.R., and Ren, B. (2014). CRISPR reveals a distal super-enhancer required for Sox2 expression in mouse embryonic stem cells. PLoS One 9, e114485.

Lim, K.H., Kim, S.R., Ramakrishna, S., and Baek, K.H. (2014). Critical lysine residues of Klf4 required for protein stabilization and degradation. Biochem Biophys Res Commun 443, 1206–1210.

Liu, Y., Beyer, A., and Aebersold, R. (2016). On the Dependency of Cellular Protein Levels on mRNA Abundance. Cell 165, 535–550.

Lu, X., Mazur, S.J., Lin, T., Appella, E., and Xu, Y. (2014). The pluripotency factor nanog promotes breast cancer tumorigenesis and metastasis. Oncogene 33, 2655–2664.

Mali, P., Yang, L., Esvelt, K.M., Aach, J., Guell, M., DiCarlo, J.E., Norville, J.E., and Church, G.M. (2013). RNA-guided human genome engineering via Cas9. Science 339, 823–826.

Mandegar, M.A., Huebsch, N., Frolov, E.B., Shin, E., Truong, A., Olvera, M.P., Chan, A.H., Miyaoka, Y., Holmes, K., Spencer, C.I., et al. (2016). CRISPR Interference Efficiently Induces Specific and Reversible Gene Silencing in Human iPSCs. Cell Stem Cell 18, 541–553.

Matsuda, T., Nakamura, T., Nakao, K., Arai, T., Katsuki, M., Heike, T., and Yokota, T. (1999). STAT3 activation is sufficient to maintain an undifferentiated state of mouse embryonic stem cells. EMBO J 18, 4261–4269.

Mlynarczyk-Evans, S., Royce-Tolland, M., Alexander, M.K., Andersen, A.A., Kalantry, S., Gribnau, J., and Panning, B. (2006). X chromosomes alternate between two states prior to random X-inactivation. PLoS Biol 4, e159.

Moorthy, S.D., Davidson, S., Shchuka, V.M., Singh, G., Malek-Gilani, N., Langroudi, L., Martchenko, A., So, V., Macpherson, N.N., and Mitchell, J.A. (2017). Enhancers and super-enhancers have an equivalent regulatory role in embryonic stem cells through regulation of single or multiple genes. Genome Res 27, 246–258.

Moorthy, S.D., and Mitchell, J.A. (2016). Generating CRISPR/Cas9 mediated monoallelic deletions to study enhancer function in mouse embryonic stem cells. J Vis Exp, e53552–e53552.

Morey, L., Santanach, A., and Di Croce, L. (2015). Pluripotency and Epigenetic Factors in Mouse Embryonic Stem Cell Fate Regulation. Mol Cell Biol 35, 2716–2728.

Nichols, J., and Smith, A. (2009). Naive and primed pluripotent states. Cell Stem Cell 4, 487–492.

Niwa, H., Burdon, T., Chambers, I., and Smith, A. (1998). Self-renewal of pluripotent embryonic stem cells is mediated via activation of STAT3. Genes Dev 12, 2048–2060.

Ohtsuka, S., Nakai-Futatsugi, Y., and Niwa, H. (2015). LIF signal in mouse embryonic stem cells. JAKSTAT 4, e1086520.

Piva, M., Domenici, G., Iriondo, O., Rabano, M., Simoes, B.M., Comaills, V., Barredo, I., Lopez-Ruiz, J.A., Zabalza, I., Kypta, R., et al. (2014). Sox2 promotes tamoxifen resistance in breast cancer cells. EMBO Mol Med 6, 66–79.

Radivojac, P., Vacic, V., Haynes, C., Cocklin, R.R., Mohan, A., Heyen, J.W., Goebl, M.G., and Iakoucheva, L.M. (2010). Identification, analysis, and prediction of protein ubiquitination sites. Proteins 78, 365–380.

Raz, R., Lee, C.K., Cannizzaro, L.A., d’Eustachio, P., and Levy, D.E. (1999). Essential role of STAT3 for embryonic stem cell pluripotency. Proc Natl Acad Sci U S A 96, 2846–2851.

Sanchez-Castillo, M., Ruau, D., Wilkinson, A.C., Ng, F.S., Hannah, R., Diamanti, E., Lombard, P., Wilson, N.K., and Gottgens, B. (2014). CODEX: a next-generation sequencing experiment database for the haematopoietic and embryonic stem cell communities. Nucleic Acids Res 43, D1117–1123.

Schoenfelder, S., Furlan-Magaril, M., Mifsud, B., Tavares-Cadete, F., Sugar, R., Javierre, B.M., Nagano, T., Katsman, Y., Sakthidevi, M., Wingett, S.W., et al. (2015). The pluripotent regulatory circuitry connecting promoters to their long-range interacting elements. Genome Res 25, 582–597.

Schuetz, A., Nana, D., Rose, C., Zocher, G., Milanovic, M., Koenigsmann, J., Blasig, R., Heinemann, U., and Carstanjen, D. (2011). The structure of the Klf4 DNA-binding domain links to self-renewal and macrophage differentiation. Cell Mol Life Sci 68, 3121–3131.

Segre, J.A., Bauer, C., and Fuchs, E. (1999). Klf4 is a transcription factor required for establishing the barrier function of the skin. Nat Genet 22, 356–360.

Sim, Y.J., Kim, M.S., Nayfeh, A., Yun, Y.J., Kim, S.J., Park, K.T., Kim, C.H., and Kim, K.S. (2017). 2i Maintains a Naive Ground State in ESCs through Two Distinct Epigenetic Mechanisms. Stem Cell Reports 8, 1312–1328.

Soheili, S., Asadi, M.H., and Farsinejad, A. (2017). Distinctive expression pattern of OCT4 variants in different types of breast cancer. Cancer Biomark 18, 69–76.

Thakore, P.I., D’Ippolito, A.M., Song, L., Safi, A., Shivakumar, N.K., Kabadi, A.M., Reddy, T.E., Crawford, G.E., and Gersbach, C.A. (2015). Highly specific epigenome editing by CRISPR-Cas9 repressors for silencing of distal regulatory elements. Nat Methods 12, 1143–1149.

Theunissen, T.W., van Oosten, A.L., Castelo-Branco, G., Hall, J., Smith, A., and Silva, J.C. (2011). Nanog overcomes reprogramming barriers and induces pluripotency in minimal conditions. Curr Biol 21, 65–71.

Tolhuis, B., Palstra, R.J., Splinter, E., Grosveld, F., and de Laat, W. (2002). Looping and interaction between hypersensitive sites in the active beta-globin locus. Mol Cell 10, 1453–1465.

Tosolini, M., and Jouneau, A. (2016). Acquiring Ground State Pluripotency: Switching Mouse Embryonic Stem Cells from Serum/LIF Medium to 2i/LIF Medium. Methods Mol Biol 1341, 41–48.

Visel, A., Blow, M.J., Li, Z., Zhang, T., Akiyama, J.A., Holt, A., Plajzer-Frick, I., Shoukry, M., Wright, C., Chen, F., et al. (2009). ChIP-seq accurately predicts tissue-specific activity of enhancers. Nature 457, 854–858.

Wei, Z., Gao, F., Kim, S., Yang, H., Lyu, J., An, W., Wang, K., and Lu, W. (2013). Klf4 organizes long-range chromosomal interactions with the oct4 locus in reprogramming and pluripotency. Cell Stem Cell 13, 36–47.

Wei, Z., Yang, Y., Zhang, P., Andrianakos, R., Hasegawa, K., Lyu, J., Chen, X., Bai, G., Liu, C., Pera, M., et al. (2009). Klf4 interacts directly with Oct4 and Sox2 to promote reprogramming. Stem Cells 27, 2969–2978.

Wray, J., Kalkan, T., and Smith, A.G. (2010). The ground state of pluripotency. Biochem Soc Trans 38, 1027–1032.

Xie, L., Torigoe, S.E., Xiao, J., Mai, D.H., Li, L., Davis, F.P., Dong, P., Marie-Nelly, H., Grimm, J., Lavis, L., et al. (2017). A dynamic interplay of enhancer elements regulates Klf4 expression in naive pluripotency. Genes Dev 31, 1795–1808.

Yeom, Y.I., Fuhrmann, G., Ovitt, C.E., Brehm, A., Ohbo, K., Gross, M., Hubner, K., and Scholer, H.R. (1996). Germline regulatory element of Oct-4 specific for the totipotent cycle of embryonal cells. Development 122, 881–894.

Zhang, P., Andrianakos, R., Yang, Y., Liu, C., and Lu, W. (2010). Kruppel-like factor 4 (Klf4) prevents embryonic stem (ES) cell differentiation by regulating Nanog gene expression. J Biol Chem 285, 9180–9189.

Zhou, B.P., Deng, J., Xia, W., Xu, J., Li, Y.M., Gunduz, M., and Hung, M.C. (2004). Dual regulation of Snail by GSK-3beta-mediated phosphorylation in control of epithelial-mesenchymal transition. Nat Cell Biol 6, 931–940.

Zhou, H.Y., Katsman, Y., Dhaliwal, N.K., Davidson, S., Macpherson, N.N., Sakthidevi, M., Collura, F., and Mitchell, J.A. (2014). A Sox2 distal enhancer cluster regulates embryonic stem cell differentiation potential. Genes Dev 28, 2699–2711.

Zuo, D., Mohr, S.E., Hu, Y., Taycher, E., Rolfs, A., Kramer, J., Williamson, J., and LaBaer, J. (2007). PlasmID: a centralized repository for plasmid clone information and distribution. Nucleic Acids Res 35, D680–684.

