## Supplemental Data for "KLF4 protein stability regulated by interaction with pluripotency transcription factors overrides transcriptional control"

**Department of Cell and Systems Biology, University of Toronto, Toronto, ON, Canada.**

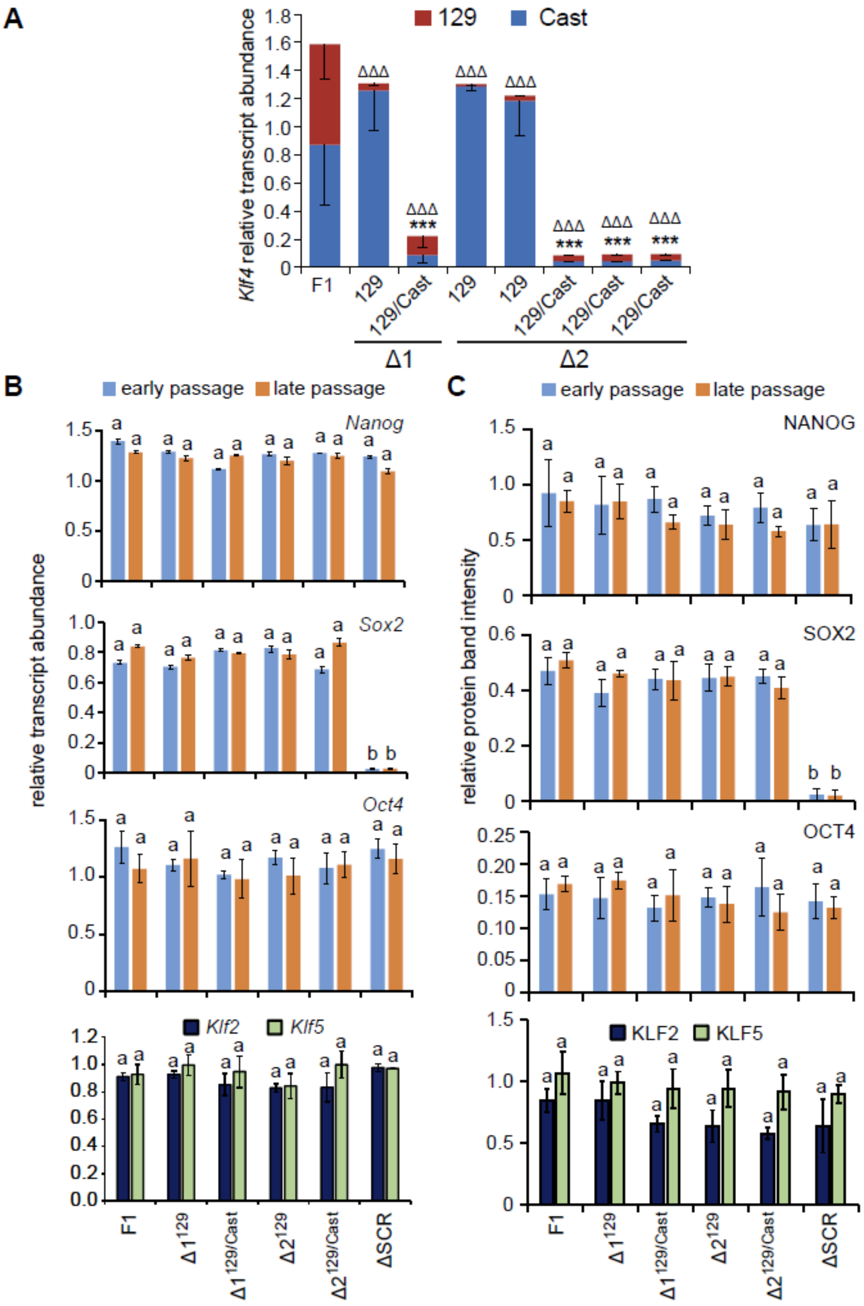


**Figure S1: The effect of CRISPR deletion of two *Klf4* enhancer regions.** A) Allele-specific primers detect *Klf4* 129 or Cast RNA in RT-qPCR from the indicated clones. Transcript levels are relative to *Gapdh*. Error bars represent standard deviation in at least three biological replicates. Statistical differences, calculated by one way ANOVA, from the F1 Cast allele are indicated by *** P < 0.001, and from the F1 129 allele by ΔΔΔ P < 0.001. B) Total transcript levels of *Oct4*, *Sox2* and *Nanog* were quantified relative to *Gapdh* from three biological replicates of both early and late passages of *Klf4* enhancer deleted clones and the $\Delta SCR^{129/Cast}$ clone (ΔSCR). *Klf2* and *Klf5* transcript levels were evaluated in late passage cells. Error bars represent standard deviation. Statistical differences determined with two way ANOVA (P < 0.05) are displayed by different letters. C) Immunoblots for OCT4, SOX2 and NANOG were quantified relative to GAPDH from three biological replicates for both early and late passages of *Klf4* enhancer deleted clones. KLF2 and KLF5 protein levels were quantified in late passage cells. Error bars represent standard deviation. Statistical differences determined by two way ANOVA (P < 0.05) are indicated by different letters.

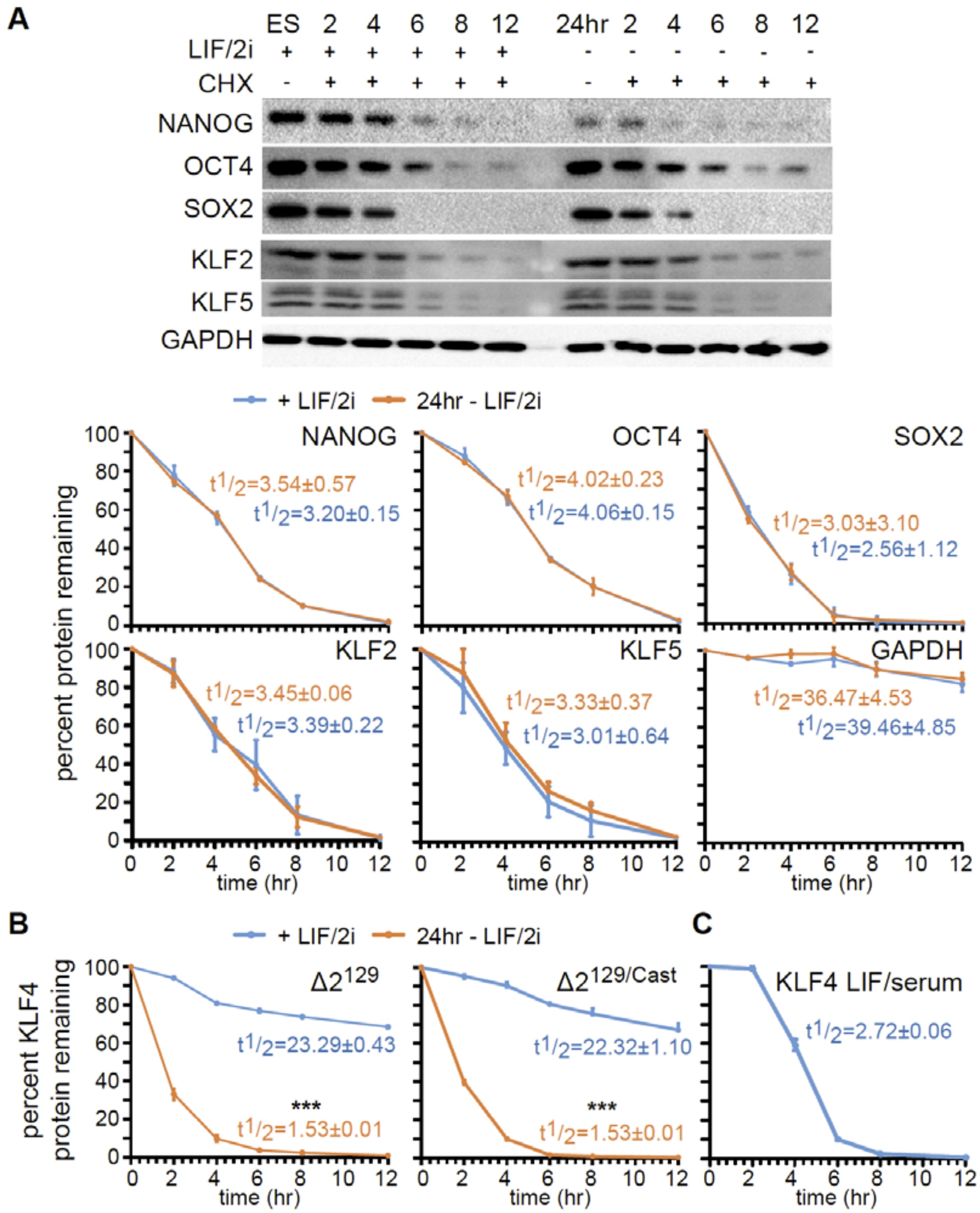


**Figure S2: OCT4, SOX2, NANOG, KLF2 and KLF5 protein stability is not affected by differentiation.** A) Immunoblots for OCT4, SOX2, NANOG, KLF2, KLF5 and GAPDH in ES cells cultured with LIF/2i and 24hr after removal of LIF/2i, sampled at 0, 2, 4, 6, 8 and 12hr after CHX treatment. GAPDH levels were used as the CHX chase assay control and displayed the expected protein half-life ($t_{½}$ >30hr). At the bottom percent remaining (OCT4, SOX2, NANOG, KLF2, KLF5 and GAPDH) protein at 0, 2, 4, 6, 8, and 12hr was calculated from the intensity of CHX treated immunoblots, measured in three biological replicates. Half-life was calculated for each time series replicate by best fit to exponential decay. Error bars represent standard deviation. Statistical differences were not identified for these proteins. B/C) Percent remaining KLF4 protein at 0, 2, 4, 6, 8, and 12hr was calculated from the intensity of CHX treated immunoblots, measured in three biological replicates for *Klf4* enhancer deleted clones maintained in LIF/2i (B) or for wild type F1 cells maintained in LIF/serum (C). Half-life was calculated for each time series replicate by best fit to exponential decay. Error bars represent standard deviation. Statistical differences determined by t test (P <0.001) are indicated as ***.

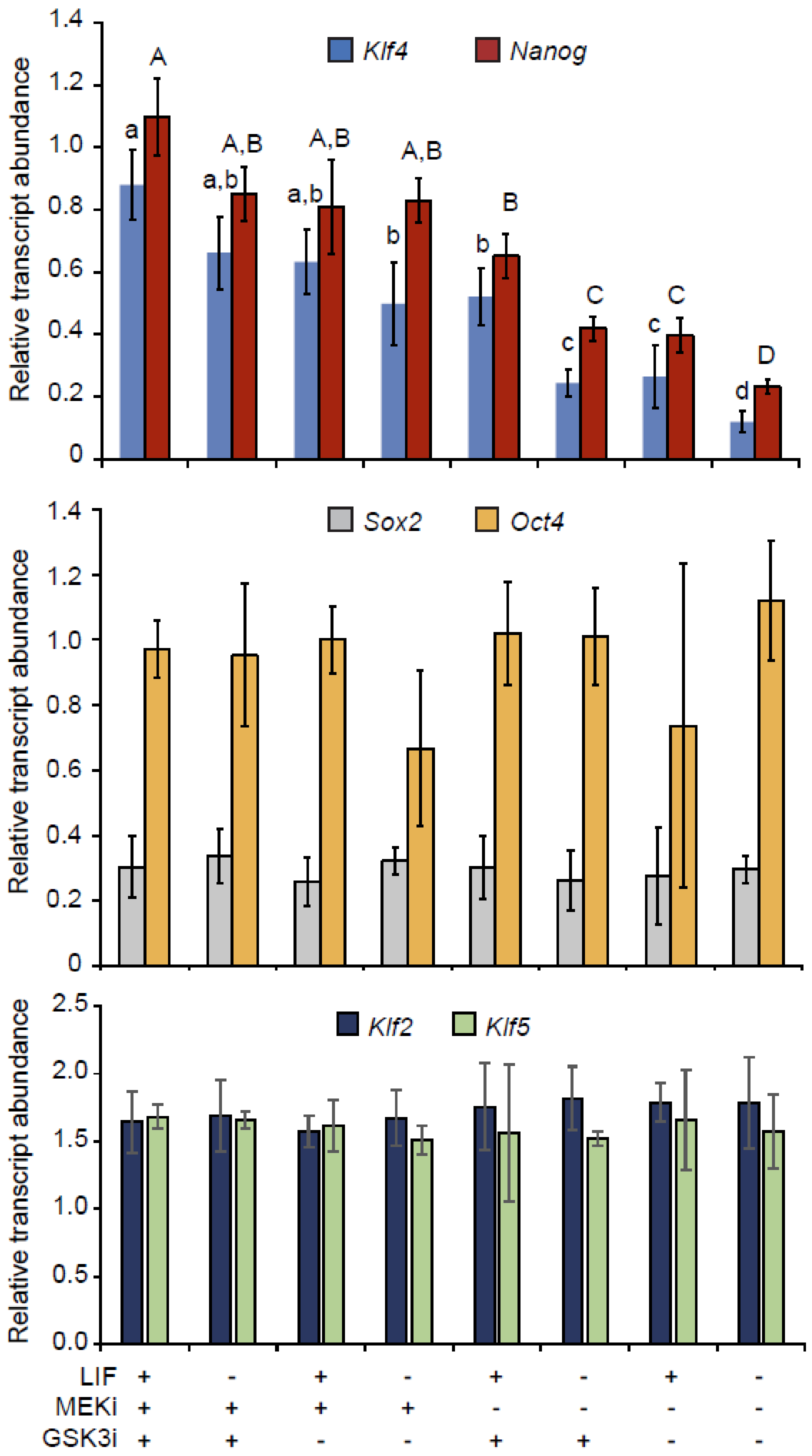


**Figure S3: Gene expression changes 12hr after removal of LIF/2i media components.** Transcript levels of *Klf4*, *Nanog*, *Sox2*, *Oct4*, *Klf2* and *Klf5* were quantified relative to *Gapdh* levels, in three biological replicates of ES cells cultured with LIF/2i and 12hr after removal of LIF/2i components. Error bars represent standard deviation. Statistical differences for each transcript were determined by one way ANOVA ($P < 0.05$) and displayed as lower case letters for *Klf4* and upper case letters for *Nanog*. No significant differences were observed for *Sox2*, *Oct4*, *Klf2* or *Klf5*.

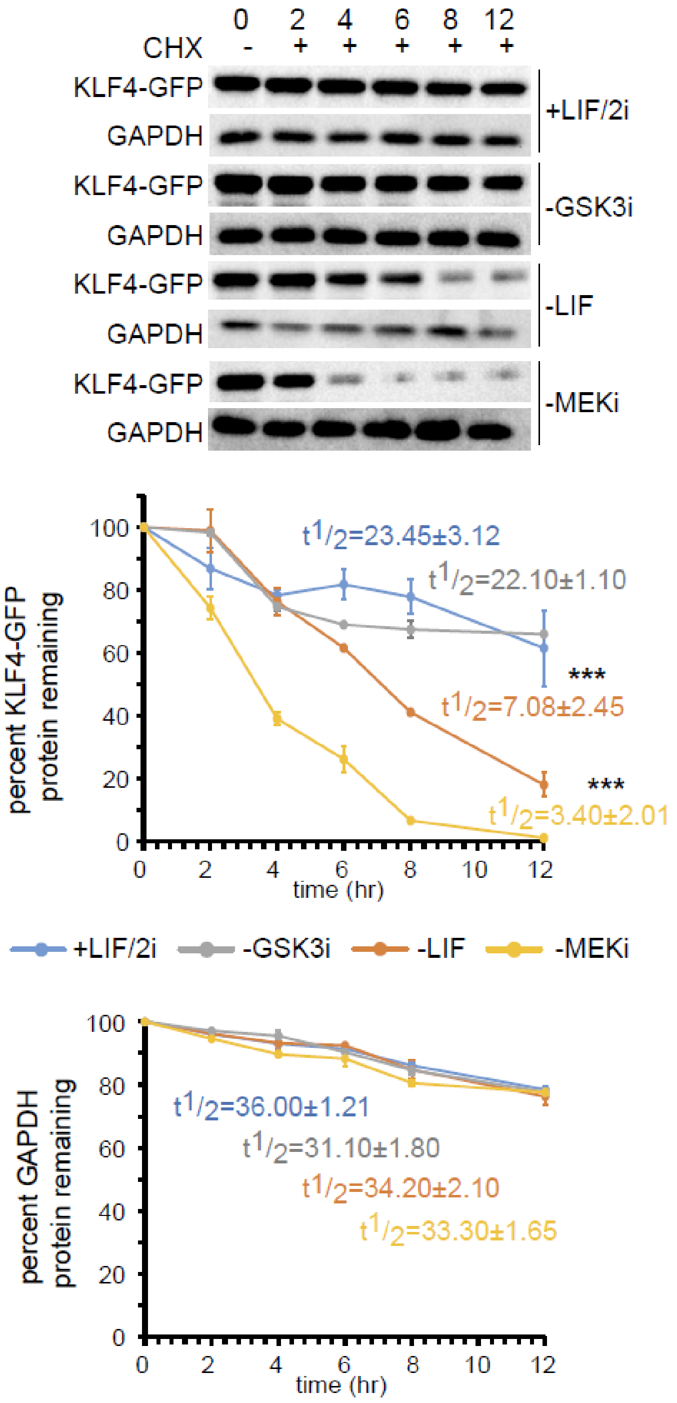


**Figure S4: KLF4-GFP protein stability is regulated by the LIF and MAPK signaling pathways.** Immunoblots for WT KLF4-GFP and GAPDH in ES cells cultured with LIF/2i and 24hr after removal of individual media components (LIF, GSK3i, and MEKi), sampled at 0, 2, 4, 6, 8 and 12hr after CHX treatment. GAPDH levels were used as the CHX chase assay control and displayed the expected protein half-life ($t_{½}$ >30hr). Percent remaining WT KLF4-GFP or GAPDH protein at 0, 2, 4, 6, 8, and 12hr was calculated from the intensity of CHX treated immunoblots, measured in three biological replicates. Half-life was calculated for each time series replicate by best fit to exponential decay. Error bars represent standard deviation. Statistical differences between protein half-life in different culture conditions compared to ES cells maintained in LIF/2i were determined by two-tailed t-test ($P < 0.001$) and are indicated by ***.

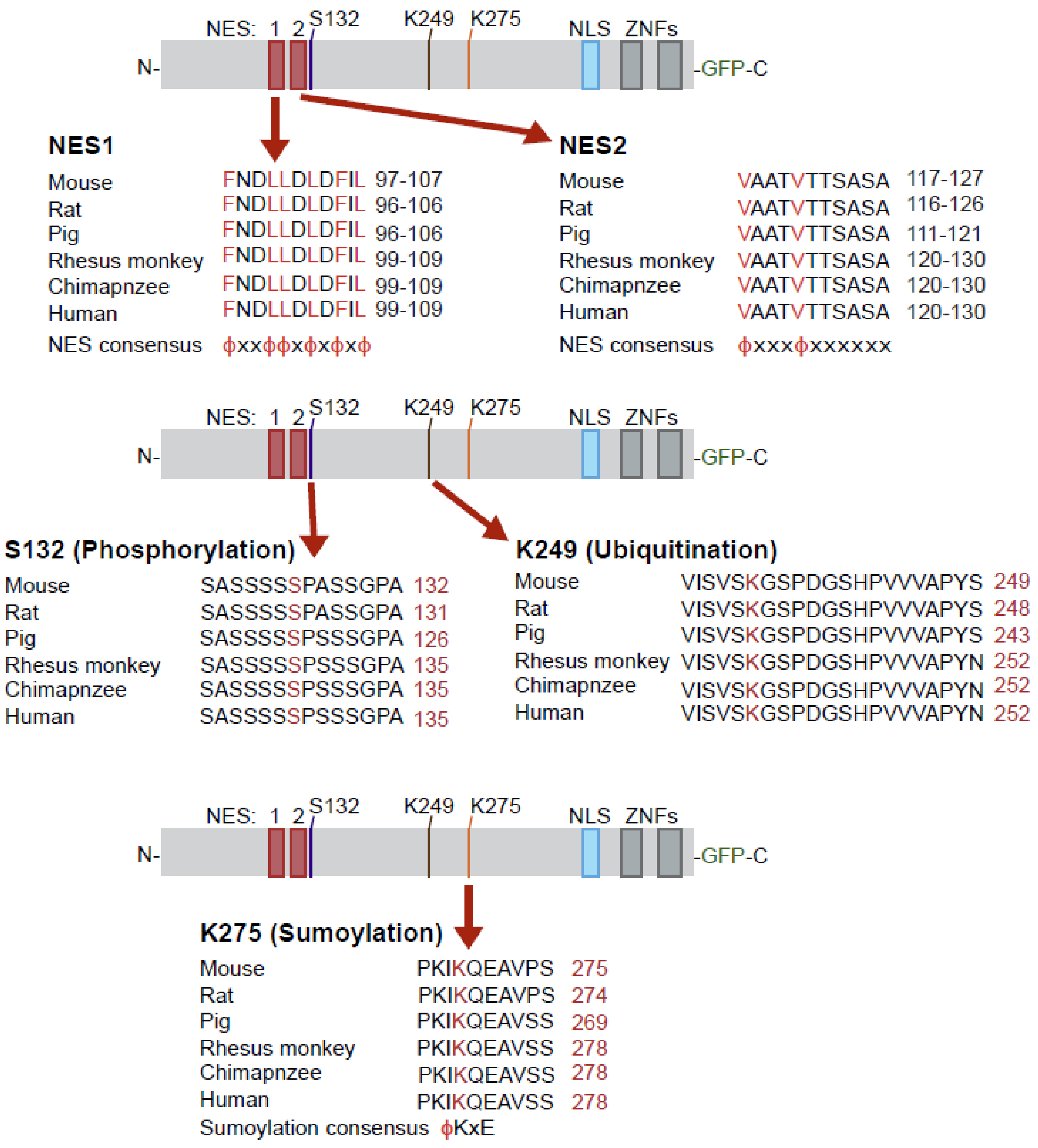


**Figure S5: KLF4 sequence conservation.** Sequence conservation for KLF4 nuclear export sequences (NES) and regions with posttranslational modifications in ES cells are indicated. The number following each sequence indicates the full range covered by the indicated sequence in each species (black) or the residue corresponding to the modified amino acid indicated in red. NLS (nuclear localization sequence), ZNFs (zinc fingers).

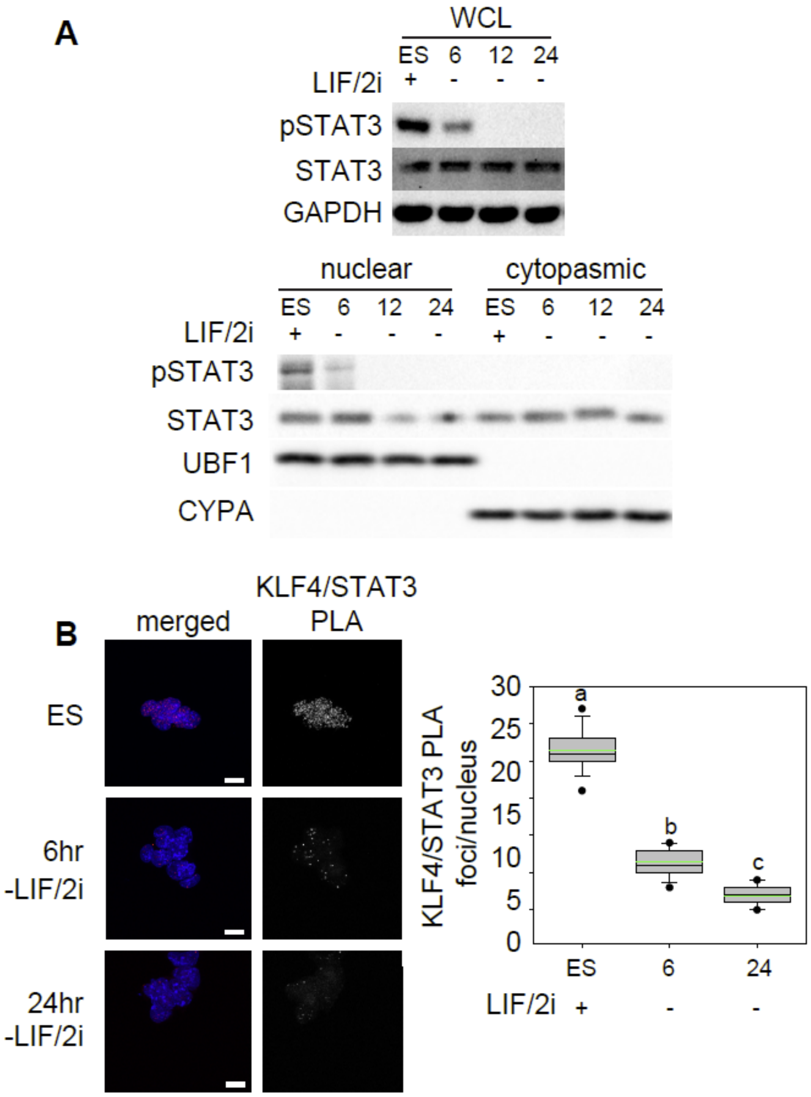


**Figure S6: Rapid changes in STAT activation and interaction with KLF4 during differentiation.** A) Immunoblots for activated pSTAT3 (Tyr705) and total STAT3. On the top whole cell lysate (WCL) prepared from ES cells cultured with LIF/2i, and 6, 12 and 24hr after removal of LIF/2i. GAPDH levels indicate equal loading. On the bottom nuclear and cytoplasmic fractions prepared from ES cells cultured with LIF/2i, 6, 12 and 24 after removal of LIF/2i were evaluated. UBF1 and CYPA were used to validate the purity of nuclear and cytoplasmic fractions respectively. B) Proximity ligation amplification (PLA) displays the interaction between KLF4/STAT3 in ES cells cultured with LIF/2i, 6 and 24hr after removal of LIF/2i. Images shown are maximum-intensity projections. Merged images display DAPI in blue and PLA in red. Scale bar = 10 µm. Box-and-whisker plots display the number of KLF4/STAT3 PLA foci per nucleus for ES cells cultured with LIF/2i, 6 and 24hr after removal of LIF/2i. Boxes indicate interquartile range of intensity values and whiskers indicate the 10th and 90th percentiles; outliers are shown as black dots. Images were collected from at least three biological replicates and ≥100 nuclei were quantified for each sample. Statistical differences between groups determined with one way ANOVA ($P < 0.05$) are indicated by different letters.

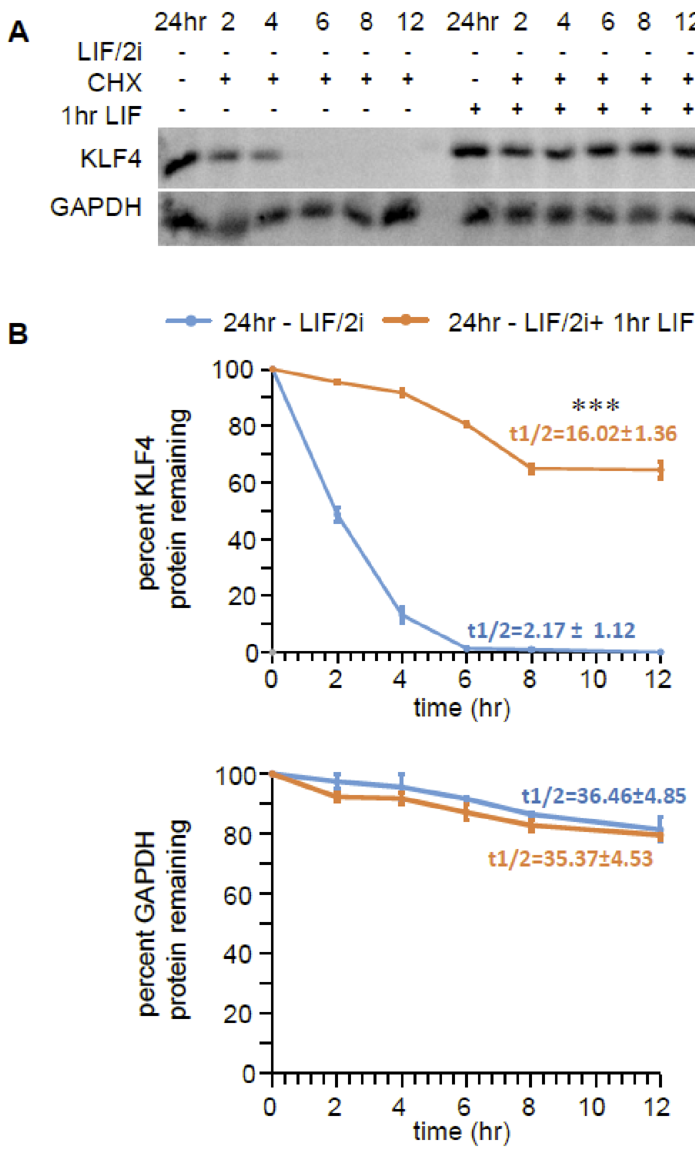


**Figure S7: LIF treatment after cyclohexamide treatment increases KLF4 stability.** A) Immunoblots for KLF4 and GAPDH in ES cells 24hr after LIF/2i removal and in cells 24hr after LIF/2i removal followed by a 1hr treatment with LIF administered at the saem time as the CHX treatment. Cells were sampled at 0, 2, 4, 6, 8 and 12 hr after CHX treatment. GAPDH levels were used as the CHX chase assay control and displayed the expected protein half-life ($t_{½}$ >30hr). B) Percent remaining KLF4 or GAPDH at 0, 2, 4, 6, 8, and 12hr was calculated from the intensity of CHX treatment immunoblots, measured in three biological replicates. Half-life was calculated for each time series replicate by best fit to exponential decay. Error bars represent standard deviation. Statistical differences between protein half-life determined by two-tailed t-test ($P < 0.001$) are indicated as ***.

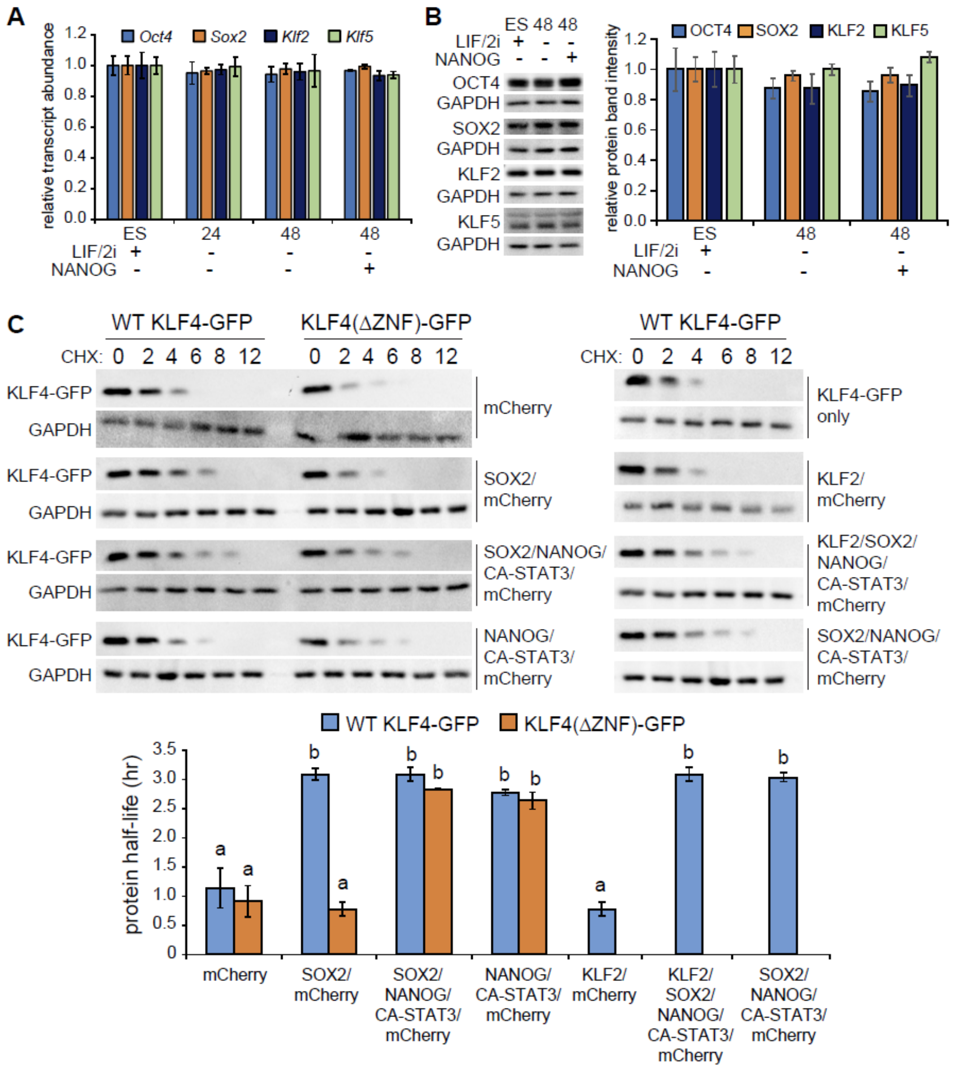


**Figure S8: KLF4 protein is stabilized by association with transcription factor complexes.** A) Transcript levels of *Oct4*, *Sox2*, *Klf2* and *Klf5* were quantified relative to *Gapdh* and undifferentiated ES cells in three biological replicates of ES cells cultured with LIF/2i (ES), 24 and 48hr after removal of LIF/2i, and in ES cells 48hr after removal of LIF/2i where Nanog-t2A-GFP was transfected 24hr after removal of LIF/2i. Error bars represent standard deviation. No significant differences were observed for *Sox2*, *Oct4*, *Klf2* or *Klf5* by one way ANOVA ($P < 0.05$). B) On the left immunoblots for OCT4, SOX2, KLF2, KLF5 and GAPDH, on the right quantification of relative protein band intensity from three biological replicates of ES cells cultured with LIF/2i (ES), 24 and 48hr after removal of LIF/2i, and in ES cells 48hr after removal of LIF/2i where Nanog-t2A-GFP was transfected 24hr after removal of LIF/2i. Error bars represent standard deviation. No significant differences were observed for OCT4, SOX2, KLF2 or KLF5 by one way ANOVA ($P < 0.05$). C) KLF4-GFP was transfected into HEK293 cells and protein stability after CHX treatment was monitored by detecting GFP and GAPDH. Cells were sampled at 0, 2, 4, 6, 8 and 12hr after CHX treatment. GAPDH levels were used as the CHX chase assay control and displayed the expected protein half-life ($t_{½}$ >30hr). Co-transfection of SOX2 but not KLF2 affected KLF4 protein stability. The greatest increase in stability was observed after co-transfection of SOX2, NANOG and constitutively active STAT3 (CA-STAT3). KLF4 containing a deletion of the zinc fingers was stabilized by the complex of SOX2, NANOG and CA-STAT3. At the bottom the calculated protein half-life is shown for the indicated conditions. Half-life was calculated for each time series replicate by best fit to exponential decay. Error bars represent standard deviation of three technical replicate immunoblots. Statistical differences determined by two-way ANOVA ($P < 0.05$) are indicated by different letters.

**Table S1: Antibody list**

| Name | Company | Catalog # | Experiment |
|---|---|---|---|
| rabbit anti-KLF4 | Abcam | AB129473 | PLA,WB, IP |
| mouse anti-KLF4 | Santa Cruz | sc-393462 | PLA, IP-WB |
| rabbit anti-NANOG | Cosmo Bio | RCAB0002P-F | PLA,WB |
| rabbit anti-NANOG | Abcam | AB80892 | PLA,WB |
| mouse anti-NANOG | BD Biosciences | 560259 | PLA,WB |
| mouse anti-SOX2 | R&D Systems | MAB2018 | PLA,WB |
| mouse anti-OCT3/4 | Santa Cruz | sc-5279 | PLA,WB |
| rabbit anti-KLF2 | Millipore | 09-820 | WB |
| rabbit anti-KLF5 | Abcam | AB137676 | WB |
| mouse anti-RNAPII-PS5 | Abcam | AB5408 | PLA,WB |
| rabbit anti-RNAPII-PS5 | Abcam | AB5131 | PLA,WB |
| mouse anti-RNAPII core (ARNA3) | Millipore Sigma | CBL221 | PLA,WB |
| mouse anti-XPO1 (CRM1) | Santa Cruz | sc-74454 | PLA, IP-WB |
| mouse anti-GFP | Origene | TA150041 | PLA |
| chicken anti-GFP | Abcam | AB13970 | WB |
| rabbit anti-CYPA | Abcam | AB131334 | WB |
| rabbit anti-UBF1 | Santa Cruz | sc-13125 | WB |
| mouse anti-GAPDH | Santa Cruz | sc-365062 | WB |
| goat anti-rabbit-HRP | Bio-Rad | 170-6515 | WB |
| goat anti-mouse-HRP | Bio-Rad | 170-6516 | WB |
| Rabbit anti ubiquitin | Abcam | AB7780 | WB |

**Table S2: List of CRISPR/Cas9 guide sequences**

| Name | Sequence |
|---|---|
| 5’Δ1 Forward | TTTCTTGGCTTTATATATCTTGTGGAAAGGACGAAACACCGCCAAGAGCGTTCGTGCCCCG |
| 5’Δ1 Reverse | GACTAGCCTTATTTTAACTTGCTATTTCTAGCTCTAAAACCGGGGCACGAACGCTCTTGGC |
| 3’Δ1 Forward | TTTCTTGGCTTTATATATCTTGTGGAAAGGACGAAACACCGAGCACAGACGGATTGAGTGA |
| 3’Δ1 Reverse | GACTAGCCTTATTTTAACTTGCTATTTCTAGCTCTAAAACTCACTCAATCCGTCTGTGCTC |
| 5’Δ2 Forward | TTTCTTGGCTTTATATATCTTGTGGAAAGGACGAAACACCGCAGATGAATTGACACGACGT |
| 5’Δ2 Reverse | GACTAGCCTTATTTTAACTTGCTATTTCTAGCTCTAAAACACGTCGTGTCAATTCATCTGC |
| 3’Δ2 Forward | TTTCTTGGCTTTATATATCTTGTGGAAAGGACGAAACACCGACTAGGGGCTCACGCGTGGT |
| 3’Δ2 Reverse | GACTAGCCTTATTTTAACTTGCTATTTCTAGCTCTAAAACACCACGCGTGAGCCCCTAGTC |

**Table S3: List of expression primers**

| Gene | Forward primer | Reverse primer | Amplicon size |
|---|---|---|---|
| *Klf4_cast* | CCCTCGTGGGAAGACAaTG | CACTACCGCAAACACACAGG | 192bp |
| *Klf4_129* | CTCGTGGGAAGACAgTGTGA | ACAGGCGAGAAACCTTACCA | 266bp |
| *Klf4* | GAAGACGAGGATGAAGCTGAC | TGGACCTAGACTTTATCCTTTCC | 94bp |
| *Nanog* | TCCCAAACAAAAGCTCTCAAG | ATCTGCTGGAGGCTGAGGTA | 165bp |
| *Sox2* | ACGCCTTCATGGTATGGTC | CGGACAAAAGTTTCCACTC | 114bp |
| *Oct4* | ATGAGGCTACAGGGACACCTT | GTGAAGTGGGGGCTTCCATA | 100bp |
| *Gapdh* | GCACCAGCATCCCTAGACC | CTTCTTGTGCAGTGCCAGGTG | 109bp |
| *Klf2* | TCATTGCAACTGGGAAGGAT | GCACAAGTGGCACTGAAAGG | 106bp |
| *Klf5* | ACGTACACCATGCCAAGTCA | GTGGGAGAGTTGGCGAATTA | 214bp |

**Table S4: List of primers for Site directed mutagenesis**

| Primer name | Sequence |
|---|---|
| K249R forward | TCGGTCATCAGTGTTAGCAGAGGAAGC |
| K249R reverse | GCTTCCTCTGCTAACACTGATGACCGA |
| K275R forward | GCATGTGCCCCAAGATTAGGCAAGAGGCGGTC |
| K275R reverse | GACCGCCTCTTGCCTAATCTTGGGGCACATGC |
| S132A forward | ccacctcggcgtcagcttcatcctcgtctgccccagcgagcagcggccctgcc |
| S132A reverse | Ggcagggccgctgctcgctggggcagacgaggatgaagctgacgccgaggtgg |
| NLS forward | cggggccacgacccgcttccgctctttggcttgg |
| NLS reverse | ccaagccaaagagcggaagcgggtcgtggccccg |
| NES1 forward | aaaggataaagtctaggtcctgttggtcgttgaactcctcggtc |
| NES1 reverse | gaccgaggagttcaacgaccaacaggacctagactttatccttt |
